# Phage-encoded single-guide RNAs subvert CRISPR-Cas9 immunity

**DOI:** 10.64898/2026.06.01.729453

**Authors:** Maximilian Feussner, Angela Migur, Jinquan Li, Stephan Riesenberg, Nils Birkholz, David Mayo-Muñoz, Holly Wakelin, Chris M. Brown, Zasha Weinberg, Rafael Pinilla-Redondo, Peter C. Fineran, Chase L. Beisel

## Abstract

Central to CRISPR technologies is the single-guide RNA (sgRNA), an engineered fusion of a processed CRISPR RNA and tracrRNA that directs Cas9 and many Cas12 nucleases to bind and cleave target DNA^1–4^. Here, we report the discovery of similarly compact viral sgRNAs (vsgRNAs) encoded by bacteriophages that counteract bacterial CRISPR-Cas9 immunity. vsgRNAs inhibit Cas9 function via two complementary routes: by sequestering the Cas9 apoenzyme, and by re-directing the Cas9 nuclease to transcriptionally silence its own promoter. vsgRNAs also can cooperate with co-encoded anti-CRISPR proteins (Acrs), including AcrIIA25.1 that blocks DNA binding by Cas9 complexed with a standard sgRNA but not with the vsgRNA. We predict that phages evolved vsgRNAs by co-opting and repurposing host-encoded long-form tracrRNAs (tracr-L) responsible for Cas9 auto-repression and a countermeasure to Acrs^5,6^. Our search also uncovered Cas9-regulating small CRISPR-associated RNAs (scaRNAs), which we predict were inserted upstream of tracrRNAs to form tracr-L but were also co-opted by phages as viral scaRNAs to suppress Cas9 immunity. Finally, we found that vsgRNAs can enable genome editing in mammalian cells, offering a natural guide RNA template for CRISPR technologies. Overall, these findings reveal that bacteriophages devised their own compact sgRNAs tailored to subvert Cas9 immunity, long preceding their rational design to program RNA-guided nucleases^2^.

## INTRODUCTION

CRISPR-Cas9 technologies have transformed genome editing and gene regulation approaches and enabled a growing set of precision gene therapies against otherwise incurable diseases^3^. Central to these biomolecular tools is the single-guide RNA (sgRNA), an ∼100-nucleotide (nt) RNA that directs Cas9 to bind and cleave DNA complementary to the guide portion of the sgRNA and flanked by a Cas9-specific protospacer-adjacent motif (PAM)^2,7^. By altering the guide sequence of the sgRNA, Cas9 can be redirected, achieving programmable and sequence-specific DNA targeting^2^.

The sgRNA comprises an engineered fusion of two RNAs principally associated with Cas9 nucleases from type II CRISPR-Cas systems, which confer adaptive immunity against bacteriophages (phages) and other mobile genetic elements (MGEs) in bacteria^1,2,8,9^. One RNA, the CRISPR RNA (crRNA), is encoded within the CRISPR array as an invader-derived spacer flanked by conserved repeats. The other RNA, the trans-activating crRNA (tracrRNA), hybridizes to a CRISPR repeat in the CRISPR array transcript, leading to processing of the RNA duplex by the endogenous endonuclease RNase III^1,4,10^. The duplex is also bound by Cas9, with the 20-24 nt processed spacer sequence guiding the nuclease to recognize and cleave the corresponding invader DNA^1,9,11^. To engineer an sgRNA, the processed crRNA-tracrRNA duplex is fused with a short stable tetraloop, eliminating the need for an additional RNA as well as RNase III. This sgRNA architecture has proven highly malleable, with further mutations and modifications improving and expanding the capabilities of CRISPR-Cas9 technologies^12–15^. The same concept has also been extended to many Cas12 nucleases that also rely on tracrRNAs for crRNA maturation^4^.

Here, we report that bacteriophages ‘invented’ their own compact sgRNA-like elements to suppress CRISPR-Cas9 immunity. These ∼90-nt viral sgRNAs (vsgRNAs) sequester the Cas9 apoenzyme to out-compete host-derived crRNAs, including those that target the phage. In addition, vsgRNAs direct Cas9 to bind, but not cleave, its own promoter through a truncated guide sequence, thereby reducing the Cas9 pool available for anti-phage defense. Beyond their inhibitory effect, vsgRNAs are frequently co-encoded with anti-CRISPR proteins that inhibit Cas9-mediated immunity without impinging on vsgRNA function. Additionally, vsgRNAs likely derived from Cas9-regulating RNAs encoded by certain type II CRISPR-Cas systems^5,16,17^, reflecting the independent co-option and short-circuiting of Cas9 self-regulatory mechanisms by phages to subvert CRISPR-Cas9 immunity. vsgRNAs thus parallel rationally-designed sgRNAs^18^ and can be applied to genome editing while also representing a distinct entry in the known catalog of protein- and RNA-based anti-CRISPR factors encoded by phages.

## RESULTS

### Discovery of a prophage-encoded sgRNA

We previously showed that phages co-opt repeats from multiple CRISPR types to sequester the corresponding Cas proteins and bypass immune defense^19^. Since searching for solitary CRISPR repeats in MGEs enabled the identification of these RNA-based anti-CRISPRs (Racrs)^19^, we asked if other CRISPR RNA components are similarly co-opted to circumvent immune defense. We focused on a long-form version of the tracrRNA, termed tracr-L, encoded within the type II-A CRISPR-Cas system in *Streptococcus pyogenes*^1^. While tracr-L hybridizes to transcribed CRISPR repeats to drive crRNA maturation similar to the standard tracrRNA, termed tracr-S, it also folds into a large sgRNA-like structure with an extended intervening loop linking a crRNA-like region with the tracrRNA (**Fig. 1a**)^5^. The ∼211-nt tracr-L in *S. pyogenes* uses its short 13-nt guide to recognize and bind the *cas9* promoter, thereby suppressing *cas9* transcription without cleaving the target DNA (**Fig. 1a**)^5^. Suppression can then be reversed to promote immunity, whether through the acquisition of new spacers that result in more crRNAs, phage-encoded anti-CRISPR proteins (Acrs) that block DNA binding by Cas9, or spontaneous mutation of the target site to boost immunity in a sub-population of cells^5,6,20^. tracr-L has been identified in 34 type II-A systems, including the one in *S. pyogenes*, but not in the other type II subtypes^5^. We therefore hypothesized that MGEs co-opted tracr-L to suppress CRISPR-Cas9 immunity.

**Figure 1.**
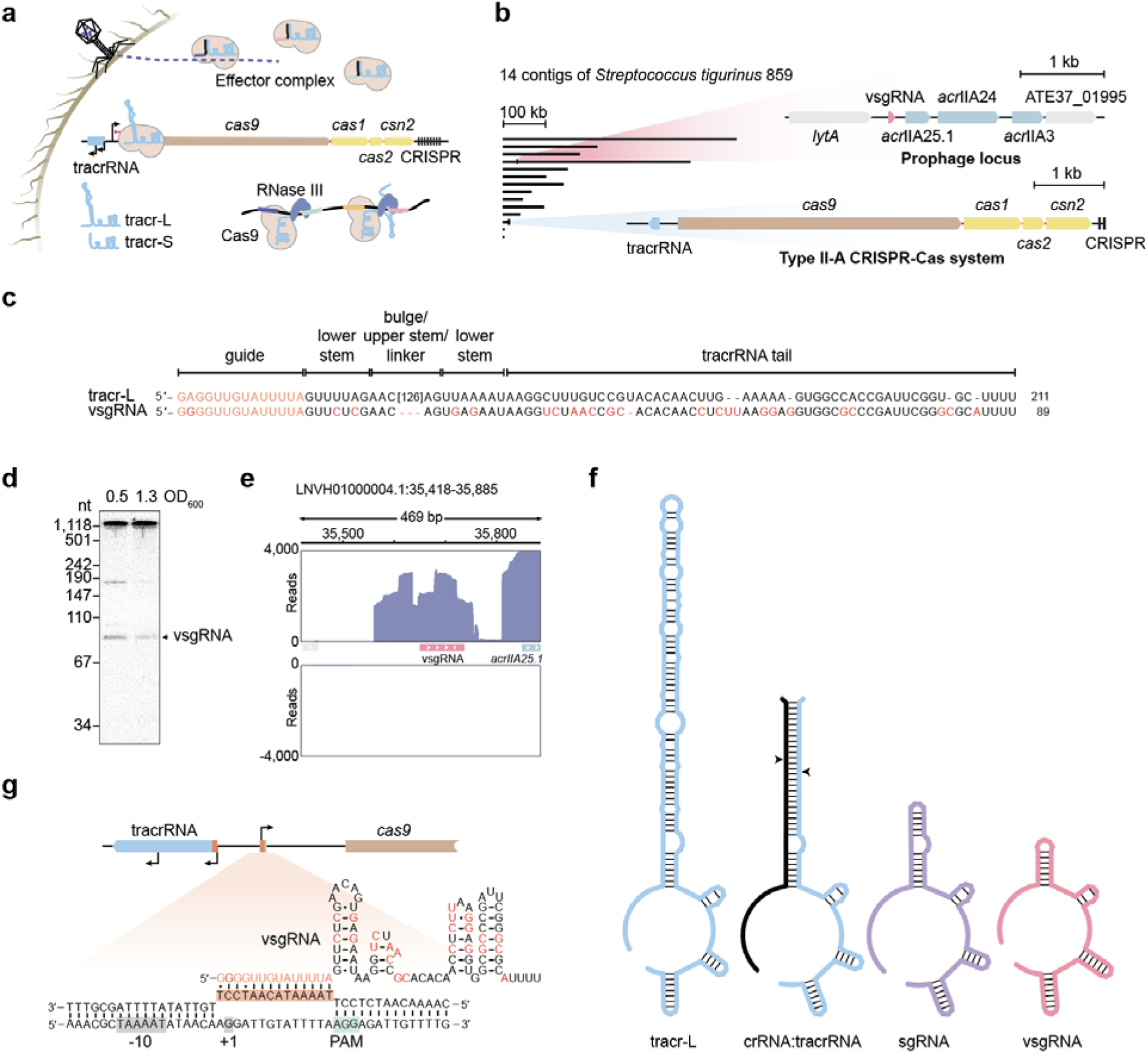
A prophage anti-CRISPR island in *S. tigurinus* expresses a compact single-guide RNA targeting the *cas9* promoter. a, Model for regulation of *cas9* transcription by the long-form tracrRNA tracr-L^5^. **b,** The 14 genomic contigs from *S. tigurinus* 859. One contig (LNVH01000012.1) encodes the type II-A CRISPR-Cas system while another (LNVH01000004.1) encodes a prophage locus with the putative vsgRNA and three Acr homologs. **c,** Sequence alignment of tracr-L and vsgRNA from *S. tigurinus*. Annotated regions relate to tracr-L. Deviations from tracr-L are marked in red, including a 126-nt region found only in the large intervening loop of tracr-L. **d,** Detection of vsgRNA expression in *S. tigurinus* at two culture turbidities (OD_600_) by northern blotting analysis. The gel image is representative of three biological replicates. **e,** RNA-seq reads from the small size fraction of total RNA of *S. tigurinus* mapped to the vsgRNA-encoding prophage locus (LNVH01000004.1). One representative biological replicate out of three is shown. **f,** Predicted secondary structures for tracr-L, the crRNA-tracrRNA duplex, sgRNA, and vsgRNA. **g,** Predicted targeting of the *cas9* promoter by the vsgRNA in *S. tigurinus*. tracr-L is contained within the tracrRNA region (blue). PAM: protospacer-adjacent motif. +1: predicted transcription start site.

An initial bioinformatics search for tracr-L homologs encoded in MGEs revealed one candidate in the bacterium *Streptococcus tigurinus* 859 (**Fig. 1b**). The candidate was found in a predicted prophage, directly upstream of three genes encoding homologs of AcrIIA25.1, AcrIIA24, and AcrIIA3^21,22^, in line with the observed clustering of anti-CRISPR genes in MGEs^19,23^. The 89-nt tracrRNA-like element aligns with the truncated guide, lower stem, and tracrRNA tail of tracr-L found in the same strain but lacks the large intervening loop (**Fig. 1c**). Northern blotting analysis and small RNA (sRNA) sequencing confirmed that the tracrRNA-like element is transcribed from the prophage, forming three transcript lengths including the shortest corresponding to the 89 nts that align with tracr-L (**Figs. 1d-e**). 5′ end mapping and structural probing of this transcript revealed a structure paralleling engineered sgRNAs, comprising a 14-nt guide, an elongated lower stem capped by a 5-nt loop, and a tracrRNA tail ending with a 3′ rho-independent terminator (**Figs. 1c,f and S1-S2**). Given the similarity of the shortest transcript to an engineered sgRNA and its presence in a putative prophage, we call this element a viral sgRNA (vsgRNA). Notably, the vsgRNA guide is predicted to target the same site as the endogenous tracr-L (**Fig. 1c**), which overlaps the transcriptional start site of the *cas9* promoter within the *S. tigurinus* genome and is flanked by a canonical 5′ NGG PAM (**Figs. 1g and S3**). Altogether, these findings reveal a prophage-encoded RNA resembling an engineered sgRNA and possessing a guide sequence corresponding to that of the Cas9-repressing tracr-L.

### The vsgRNA sequesters Cas9 protein

The similarity between the identified vsgRNA and an engineered sgRNA (**Fig. S4a**) suggests that the vsgRNA can bind Cas9, potentially preventing the nuclease from utilizing invader-targeting crRNAs in a mechanism similar to Racrs^5,24^. To first assess whether the vsgRNA can bind Cas9, we used the *Streptococcus pyogenes* Cas9 (SpCas9) as a convenient model (63% aa similarity to the Cas9 in *S. tigurinus* (StCas9) and the same 5′ NGG PAM, **Figs. S3 and S5**)^2,25^. SpCas9 bound the vsgRNA ∼9-fold tighter than an equivalent sgRNA *in vitro* (**Figs. 2a and S6**), prompting us to test whether vsgRNA binding interferes with Cas9-mediated immunity. To remove the confounding effects of the co-encoded Acrs as well as targeting of the *cas9* promoter, we expressed StCas9, a phage-targeting sgRNA, and the vsgRNA in *Escherichia coli*, using a promoter unrelated to the native *Stcas9* promoter to drive nuclease expression. Phage T7 infection was reduced by ∼10^4^-fold by StCas9 with a sgRNA targeting the *gp7* gene relative to a non-targeting guide (**Fig. 2b**). In contrast, co-expression of the vsgRNA abolished phage targeting by StCas9 with the sgRNA and allowed T7 propagation. Similar inhibition by the vsgRNA was observed when using SpCas9 in place of StCas9 (**Fig. 2b**). Together, these results show that the vsgRNA can inhibit anti-phage defense by binding and sequestering available Cas9 molecules.

**Figure 2.**
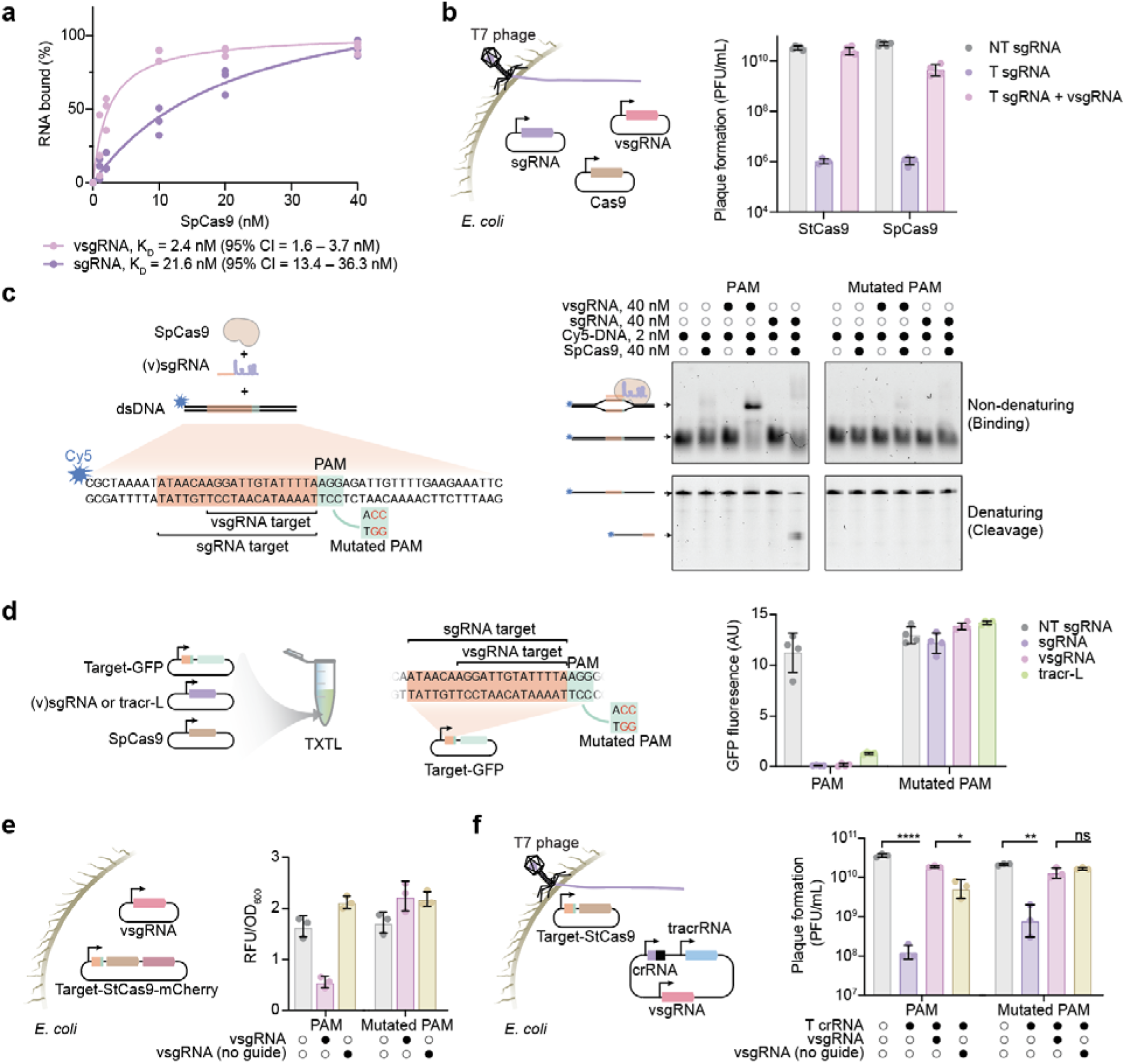
The *S. tigurinus* vsgRNA enacts layered inhibition of CRISPR-Cas9 immunity. a, Measured binding affinity of the *S. tigurinus* vsgRNA and equivalent sgRNA to SpCas9 *in vitro* using an electrophoretic mobility shift assay (EMSA). Individual EMSAs assessing vsgRNA and sgRNA binding to Cas9 are shown in **Figure S6**. Dots represent individual replicates. K_D_ values represent the mean and 95% confidence interval for three independent replicates. **b,** vsgRNA- mediated inhibition of StCas9 and SpCas9 immunity against infection by T7 phage. T, targeting. NT, non-targeting. Circles represent individual measurements. Bars and error bars represent the geometric mean and geometric standard deviation from three to six biological replicates. Cas9 was expressed from a constitutive promoter lacking the vsgRNA target site. **c,** Assessing Cas9-mediated DNA binding and cleavage by the vsgRNA *in vitro*. Left: overview of the combined components. Right: resolved labeled DNA under non-denaturing (top) and denaturing (bottom) conditions and imaged for Cy5. Gel images are representative of three independent experiments. **d,** PAM-dependent silencing of a GFP reporter by the vsgRNA in TXTL. Left: overview of the combined components in a TXTL reaction. Middle: sequence of the target site and PAM. Right: GFP fluorescence following DNA targeting with different guide RNAs. NT, non-targeting. Circles represent individual measurements. Bars and error bars represent the mean and standard deviation from four independent measurements. **e,** PAM-dependent silencing of an mCherry reporter by the vsgRNA in *E. coli*. Cas9 and mCherry were expressed from a constitutive promoter, followed by the vsgRNA target site. Separate RBS sequences were cloned upstream of each ORF. Circles represent individual measurements. Bars and error bars represent the geometric mean and geometric standard deviation from three biological replicates. RFU: Relative Fluorescence Units. **f**, Impact of Cas9 auto-repression with the vsgRNA on Cas9-mediated immunity against infection by T7 phage. T, targeting. Cas9 was expressed from a constitutive promoter followed by the vsgRNA target site. Circles represent individual measurements. Bars and error bars represent the geometric mean and geometric standard deviation from three biological replicates. *, p < 0.05. **, p < 0.01. ***, p < 0.001. ****, p < 0.0001 based on an unpaired t-test on geometric means with two tails.

### The vsgRNA induces Cas9 auto-repression

Beyond sequestering Cas9, the vsgRNA has the potential to auto-repress *cas9* transcription by targeting its promoter, analogous to tracr-L. Probing this possibility *in vitro*, we found that the vsgRNA directs SpCas9 to bind, but not cleave, the PAM-flanked DNA target sequence present in the *cas9* promoter, while a standard sgRNA led to binding and cleavage of the same site (**Figs. 2c and S4**). To interrogate whether vsgRNA-mediated binding of SpCas9 to this site blocks gene expression, we applied cell-free transcription-translation (TXTL)^26^. When placing the PAM-flanked target site immediately downstream of a constitutive promoter, the vsgRNA yielded 99% repression of a GFP reporter compared with a non-targeting control, while tracr-L yielded 89% repression (**Fig. 2d**). Furthermore, mutating the PAM in the target site abolished transcriptional repression (**Fig. 2d**). Gene repression by the vsgRNA was also observed *in vivo*. By using a transcriptionally fused mCherry reporter, the vsgRNA and StCas9 elicited a 3.3-fold gene repression in *E. coli*, which was abolished upon mutating the PAM or removing the vsgRNA guide sequence (**Fig. 2e**). Repression was dose-dependent, as reporter silencing improved with vsgRNA promoter strength (**Fig. S7a**). Thus, the vsgRNA can guide Cas9 to bind the predicted DNA target and block transcription.

Using the same promoter configuration to express Cas9, we evaluated the extent to which Cas9 auto-repression could enhance immune suppression over Cas9 sequestration alone. When infecting with T7 phage, combining both mechanisms led to 3.3-fold more plaques than sequestration alone by the vsgRNA without a guide (**Fig. 2f**). Moreover, using a PAM mutant that eliminates Cas9 auto-repression, plaque formation was unaffected by the vsgRNA having or lacking the guide (**Fig. 2f**). Both mechanisms were also dependent on vsgRNA expression, with stronger promoters leading to greater plaque formation (**Fig. S7b**). Thus, vsgRNA-directed Cas9 auto-repression contributes to immune suppression, even though Cas9 sequestration was the primary contributor. Altogether, the vsgRNA can subvert Cas9-mediated phage defense by both sequestering Cas9 protein and reducing Cas9 expression.

### Co-encoded Acrs do not inhibit vsgRNA function

The vsgRNA in *S. tigurinus* is encoded upstream of three Acrs (AcrIIA25.1, AcrIIA24, and AcrIIA3), with all three expressed and Cas9 expression silenced based on RNA-seq analysis (**Fig. S8**). Although Acrs exhibit a range of inhibitory mechanisms^27^, we asked whether the co-encoded Acrs would interfere with the functions of the vsgRNA. If anything, this scenario was possible given that AcrIIA25.1 blocks DNA binding by interacting with the Cas9-sgRNA complex^21,28^, AcrIIA24 blocks DNA cleavage but not DNA binding^21^, and AcrIIA3 remains uncharacterized^22^ (**Fig. 3a**).

**Figure 3.**
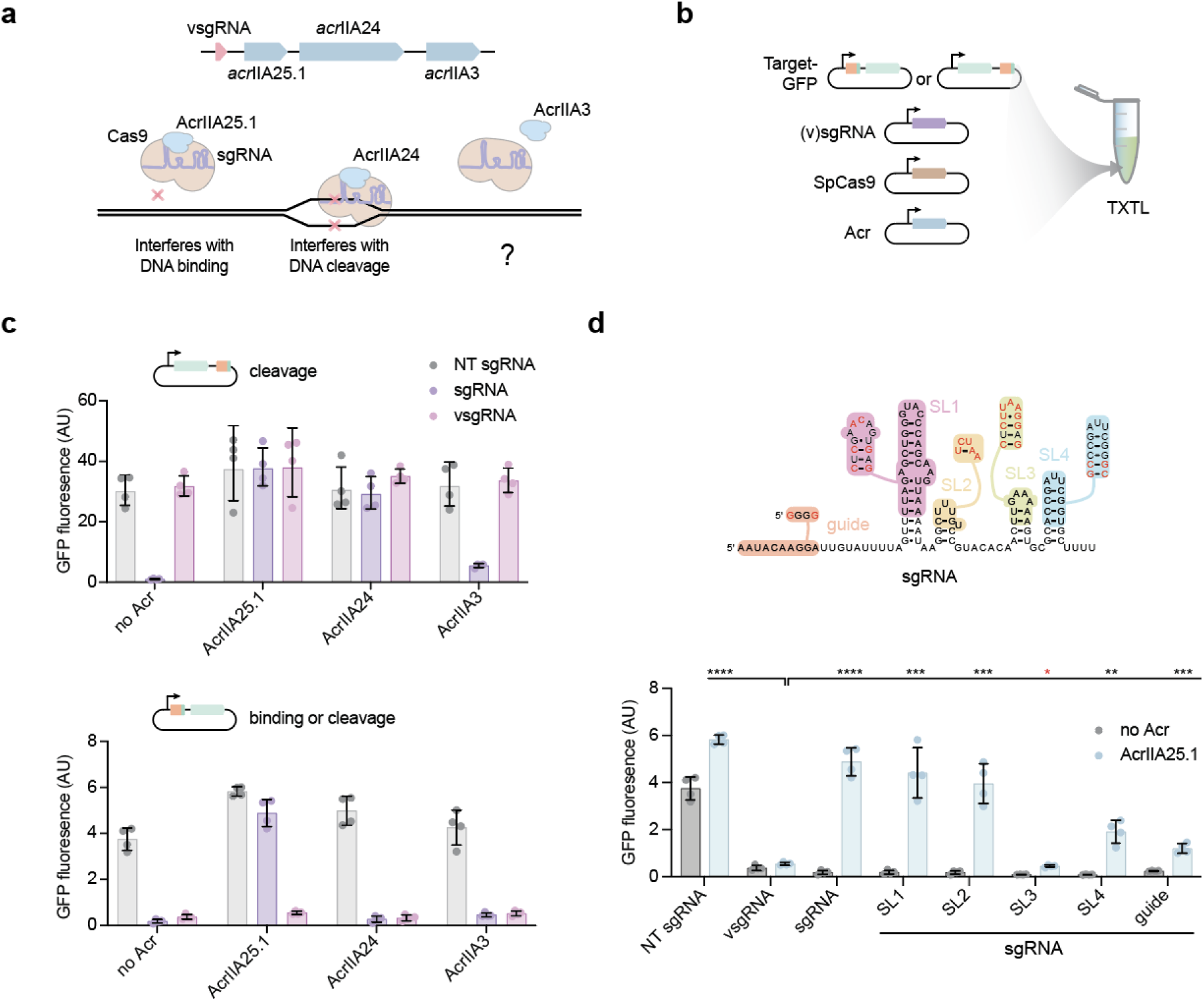
The *S. tigurinus* vsgRNA cooperates with co-encoded anti-CRISPR proteins to suppress Cas9 immunity. a, Known inhibitory mechanisms of the co-encoded Acrs. **b,** Components added to the TXTL reaction in c and d. The target sequence is the same as in Figure 2d. **c,** Impact of Acrs on GFP reporter silencing via Cas9-mediated binding (bottom) or cleavage (top). Circles represent individual measurements. Bars and error bars represent the mean and standard deviation from four independent measurements. **d,** Impact of swapping vsgRNA domains into the sgRNA on inhibition by AcrIIA25.1. Top: swapped domains. Bottom: GFP fluorescence following DNA targeting with different guide RNAs. Circles represent individual measurements. Bars and error bars represent the mean and standard deviation from four independent measurements. Statistical comparisons are between the vsgRNA and individual sgRNA variants in the presence of AcrIIA25.1. *, p < 0.05. **, p < 0.01. ***, p < 0.001. ****, p < 0.0001 based on an unpaired t-test with two tails. For the red asterisk, the mean for SL3 is below the mean of the vsgRNA. NT, non-targeting.

We began by using TXTL to interrogate how each Acr acts on Cas9 (**Fig. 3b**). Each Acr fully or modestly inhibited sgRNA-guided Cas9 cleavage of the GFP reporter plasmid backbone (**Fig. 3c**, top), where cleavage of the DNA target downstream of the reporter construct would lead to rapid plasmid degradation through RecBCD^26,29^. To gauge whether each Acr could inhibit DNA binding by Cas9, we encoded the same target in the 5′ untranslated region (UTR) of the GFP reporter gene^26^ (**Fig. 3c**, bottom). For this configuration, AcrIIA24 and AcrIIA3 failed to disrupt reporter silencing, in line with AcrIIA24 blocking DNA cleavage but not binding^21^ and suggesting that AcrIIA3 functions in a similar capacity (**Fig. 3c**, bottom). While AcrIIA25.1 disrupted reporter silencing by the sgRNA in line with its known inhibition of DNA binding^21,28^, it did not disrupt reporter silencing by the vsgRNA (**Fig. 3c**). Altogether, the three co-encoded Acrs do not interfere with Cas9 binding and gene repression by the vsgRNA in *S. tigurinus*.

The ability of AcrIIA25.1 to block DNA binding by Cas9 complexed with the sgRNA but not the vsgRNA was surprising given that the vsgRNA and the sgRNA share 55% nt identity and the same general secondary structure. We therefore sought to identify which sequences unique to the vsgRNA allow it to escape inhibition by AcrIIA25.1. Replacing different regions of the sgRNA with those of the vsgRNA demonstrated that stem-loop 3 was the only portion able to completely reverse inhibition by AcrIIA25.1 (**Fig. 3d**). As support, a published cryoEM structure of AcrIIA25.1 bound to the Cas9-sgRNA complex shows direct contacts between this stem-loop and the Acr^28^. Replacing other regions of the sgRNA with those from the vsgRNA also resulted in measurable, albeit smaller, reversals (**Fig. 3d**), suggesting that other features unique to the vsgRNA mitigate inhibition by AcrIIA25.1. In total, vsgRNA complements its three co-encoded Acrs in *S. tigurinus*, either because the Acrs specifically block DNA cleavage or are ineffective against Cas9 bound to the vsgRNA.

### vsgRNAs are exapted from long-form tracrRNAs

Validating the vsgRNA in *S. tigurinus* as a Cas9 inhibitor led us to examine the distribution and origins of vsgRNAs. Given the close resemblance between the *S. tigurinus* vsgRNA and the equivalent sgRNA derived from the endogenous II-A system, we built co-variance models capturing expected nucleotide frequencies in an RNA with a conserved secondary structure using 79 previously characterized sgRNAs^30^ and one generated sgRNA covering all three subtypes of type II CRISPR-Cas systems (**Table S1**). The covariance models were then utilized to identify sgRNA-like elements in bacterial and phage genomes (**Table S2**). The search revealed 170 such elements associated with II-A and II-C systems and 18 of the 80 sgRNA groups (**Fig. 4a and Table S2**). Furthermore, 94% appear within putative phage or prophage regions (**Fig. 4b and Table S2**), while ∼80% associate with characterized or predicted Acrs or verified Acr-associated proteins (Acas) (**Fig. 4b and Table S2**). Among these Acrs, many would be expected to not inhibit their co-encoded vsgRNAs based on their known mechanisms of action, in line with our previous insights into the three Acrs co-encoded the *S. tigurinus* vsgRNA (**Fig. 3**). For example, CasPRs block transcription by binding within the *cas* coding regions^31^, AcrIIA5 inhibits DNA cleavage^32^, and AcrIIA18 truncates the crRNA guide to 15 nts^33^. In contrast, some Acrs, such as AcrIIA27 that inhibits DNA binding^34^, could interfere with gene repression by the vsgRNA. However, this mode of inhibition would still allow the vsgRNA to sequester Cas9 that dominated in our experiments (**Fig. 2b,f**). Several vsgRNAs were also co-encoded with AcrVIB1, an anti-CRISPR of the type VI-B1 system^35,36^, which can co-occur and cooperate with type II systems^37^. To directly explore if these predicted vsgRNAs can inhibit Cas9 immunity, we tested seven representative examples from different *Streptococcus* strains in addition to the *S. tigurinus* vsgRNA (**Figs. 4c and S9, Table S3**). When targeting T7 phage with SpCas9 or StCas9 expressed from a heterologous promoter in *E. coli*, each of the seven candidates recovered plaque formation for both nucleases (**Fig. 4c**), demonstrating their function as vsgRNAs that can sequester Cas9 to block anti-phage immunity.

**Figure 4.**
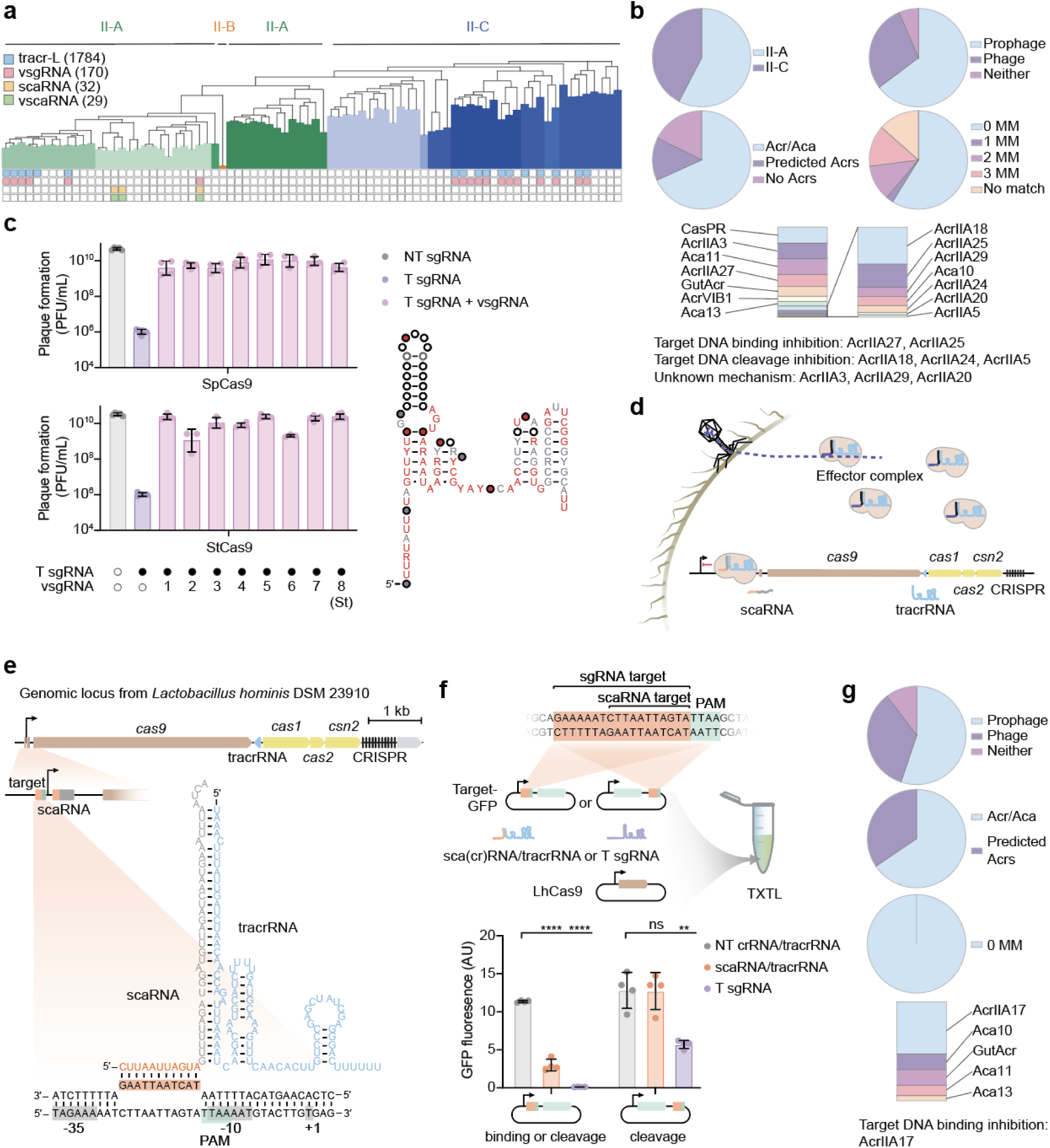
Cas9-regulating RNAs gave rise to viral sgRNAs and scaRNAs. a, Presence of Cas9-regulating RNAs and inhibitory RNAs predicted based on similarity with each of the 80 sgRNAs. sgRNAs were clustered and sorted based on their sequences and structures. The number of each identified Cas9-regulating RNAs and their viral counterparts are indicated in parentheses. **b,** Properties of the identified 170 vsgRNAs. Upper left: association with Cas9 subtypes. Upper right: association with (pro)phage regions. Lower left: association with predicted or verified Acrs or Acas. Lower right: number of mismatches (MM) between tracr-L and vsgRNA guide sequences in the same sgRNA group. No match is defined as any pairing with >3 mismatches. Bottom: associated Acrs or Acas and their known mechanisms of action. **c,** Inhibition of SpCas9 and StCas9 immunity by eight vsgRNAs from *Streptococcus* species including *S. tigurinus* 859 (vsgRNA_8). Left: efficiency of plating following infection by T7 phage targeted by SpCas9 or StCas9. T, targeting. NT, non-targeting. See **Table S3** for the list of vsgRNAs and **Figure S9** for their structures. Circles represent individual measurements. Bars and error bars represent the geometric mean and geometric standard deviation from three to six biological replicates. Cas9 was expressed from a constitutive promoter lacking the vsgRNA target site. Right: consensus vsgRNA sequence. Solid circles represent the frequency of a nucleotide being present at a specific alignment position, while the colored nucleotides denote the frequency of a specific nucleotide identity across the aligned sequences. Thresholds for 97% (red), 90% (black), and 75% (grey) are shown. **d,** Regulation of Cas9 transcription by a scaRNA. **e,** Location and promoter targeting by the Cas9-regulating scaRNA in *Lactobacillus hominis* DSM 23910. Orange boxes indicate the scaRNA guide and DNA target. The PAM sequence was predicted with Protein2PAM^91^. **f,** Gene repression by the Cas9-regulating scaRNA. Top: overview of the combined components in a TXTL reaction. Bottom: GFP fluorescence following DNA targeting with different guide RNAs. NT, non-targeting. Circles represent individual measurements. Bars and error bars represent the mean and standard deviation from four independent measurements. *, p < 0.05. **, p < 0.01. ***, p < 0.001. ****, p < 0.0001 based on an unpaired t-test with two tails. Statistical comparisons are between the NT crRNA/tracrRNA and either of the other RNA pairs with the same target site location. **g,** Properties of the identified 29 vscaRNAs. Top: association with (pro)phage regions. Middle: association with predicted or verified Acrs or Acas. Bottom: number of mismatches (MM) between scaRNA and vscaRNA guide sequences in the same gRNA group, with associated Acrs or Acas and their known mechanisms of action.

Given that the *S. tigurinus* vsgRNA was identified in part based on its homology to tracr-L in the same strain, we explored the extent to which predicted vsgRNAs have similar sequences to tracr-L and even share guide sequences. We first used the covariance models to search for additional tracr-L sequences, which expanded the known tracr-L catalog by ∼50-fold relative to prior work and uncovered their presence in type II-C systems (**Figs. 4a-b and Table S4**)^5^. In this expanded dataset, 15 of the 18 sgRNA groups with a putative vsgRNA also contained a tracr-L (**Fig. 4a**). Importantly, of these vsgRNAs, 59% shared the exact guide sequence as a tracr-L guide, while an additional 28% contained up to three mismatches (**Fig. 4b**). The remaining putative vsgRNAs contained guides with no obvious DNA target in the immediate vicinity of the *cas9* promoter (**Fig. 4a and Table S2**), suggesting that these vsgRNAs principally operate through Cas9 sequestration or may have other unknown targets. The frequent co-occurrence of vsgRNAs with closely related tracr-L variants across taxa suggests that their origin is most parsimoniously explained by the repeated, independent co-option of host tracr-L by phages for Cas9 sequestration and auto-repression, followed by subsequent modifications to prevent functional interference by co-encoded Acr proteins.

### Discovery of Cas9-regulating scaRNAs and exapted viral scaRNAs

While the vast majority of the putative vsgRNAs matched identified tracr-L sequences in the same sgRNA-derived group, one group contained 18 putative vsgRNAs but no tracr-L (**Fig. 4a**, column with red square but no blue square). This anomaly prompted us to explore whether an alternative mode of Cas9 inhibition encoded within the CRISPR-Cas system might operate in this lineage that could have given rise to this group of vsgRNAs. Searching the *Gemella morbillorum* genome harboring a vsgRNA for other copies of its guide sequence revealed an exact match downstream of *cas9* and the tracrRNA, which was flanked by a sequence with extensive complementarity to the tracrRNA anti-repeat region (**Figs. S10a-c**). This element is reminiscent of the small CRISPR-associated RNA (scaRNA), an RNA expressed from the type II-B CRISPR-Cas system in *Francisella novicida* that mimics a CRISPR guide-repeat and is encoded upstream of the CRISPR array^16^. Prior work showed that the scaRNA hybridizes to the tracrRNA and directs Cas9 to suppress the expression of virulence genes through a truncated guide (**Fig. 4d**)^16,17,38^. The element in *G. morbillorum* could act similarly but target the *cas9* promoter in the −35 element (**Fig. S10c**). Searching for similar elements that target the *cas9* promoter and are encoded within or around the CRISPR-Cas system revealed a total of 32 within three of the sgRNA groups from the co-variance models (**Fig. 4a and Table S5**). These putative Cas9-regulating scaRNAs are present either downstream of the CRISPR array or within the predicted 5′ UTR of the *cas9* gene (**Figs. 4e and S11**). In addition, tracr-L could be interpreted as a Cas9-regulating scaRNA upstream of the tracrRNA, suggesting that tracr-L evolved from a scaRNA insertion event.

To validate these elements as Cas9-regulating scaRNAs, we chose one representative example from the II-A CRISPR-Cas system in the commercially available bacterium *Lactobacillus hominis* (**Fig. 4e**). Small RNA-seq confirmed expression of the scaRNA upstream of the gene encoding the *L. hominis* Cas9 (LhCas9) (**Fig. S12**). In TXTL, combining expressed LhCas9 with *in vitro*-transcribed and annealed scaRNA and tracrRNA led to significant repression of the GFP reporter when the target was placed in the 5′ UTR but not in the plasmid backbone (**Fig. 4f**). This scaRNA-mediated gene repression without DNA cleavage parallels our previous observations with the vsgRNA (**Fig. 3c**). Thus, type II CRISPR-Cas systems encode scaRNAs that can transcriptionally repress Cas9 expression, presumably providing a negative feedback loop similar to tracr-L^5^.

The discovery of Cas9-regulating scaRNAs opened the possibility that phages directly co-opted these RNAs as viral scaRNAs (vscaRNAs). We initially searched for elements containing a similar guide and repeat sequence present in the same genome and then used matches to identify homologs in other genomes. This search revealed 29 scaRNA-like elements in three of the sgRNA groups from the co-variance models containing at least one Cas9-regulating scaRNA (**Fig. 4a and Table S6**), with the scaRNAs and vscaRNAs sharing the same guide sequence targeting the *cas9* promoter. Furthermore, 90% are encoded by predicted phages or prophages, while all are located nearby to an Acr or Aca and perfectly match the scaRNA guide within the same sgRNA co-variance model (**Fig. 4g and Table S6**). We tested the vscaRNA associated with the *L. hominis* scaRNA and an equivalent crRNA and tracrRNA in TXTL (**Figs. S13a-b**). The vscaRNA enacted targeted GFP repression only when the guide was lengthened to 20 nts. However, we attribute the lack of repression by the unaltered vscaRNA to a large bulge only present in the lower stem of the vscaRNA-tracrRNA duplex – a distinction absent in other crRNA-tracrRNA and vscaRNA-tracrRNA pairs (**Figs. S13a-b and S14**). Altogether, these results lend evidence to a model in which phages co-opted two distinct classes of Cas9-regulating RNAs (i.e., tracr-L and scaRNA) to short-circuit regulation and suppress Cas9 immunity (**Figs. 5a-b**).

**Figure 5.**
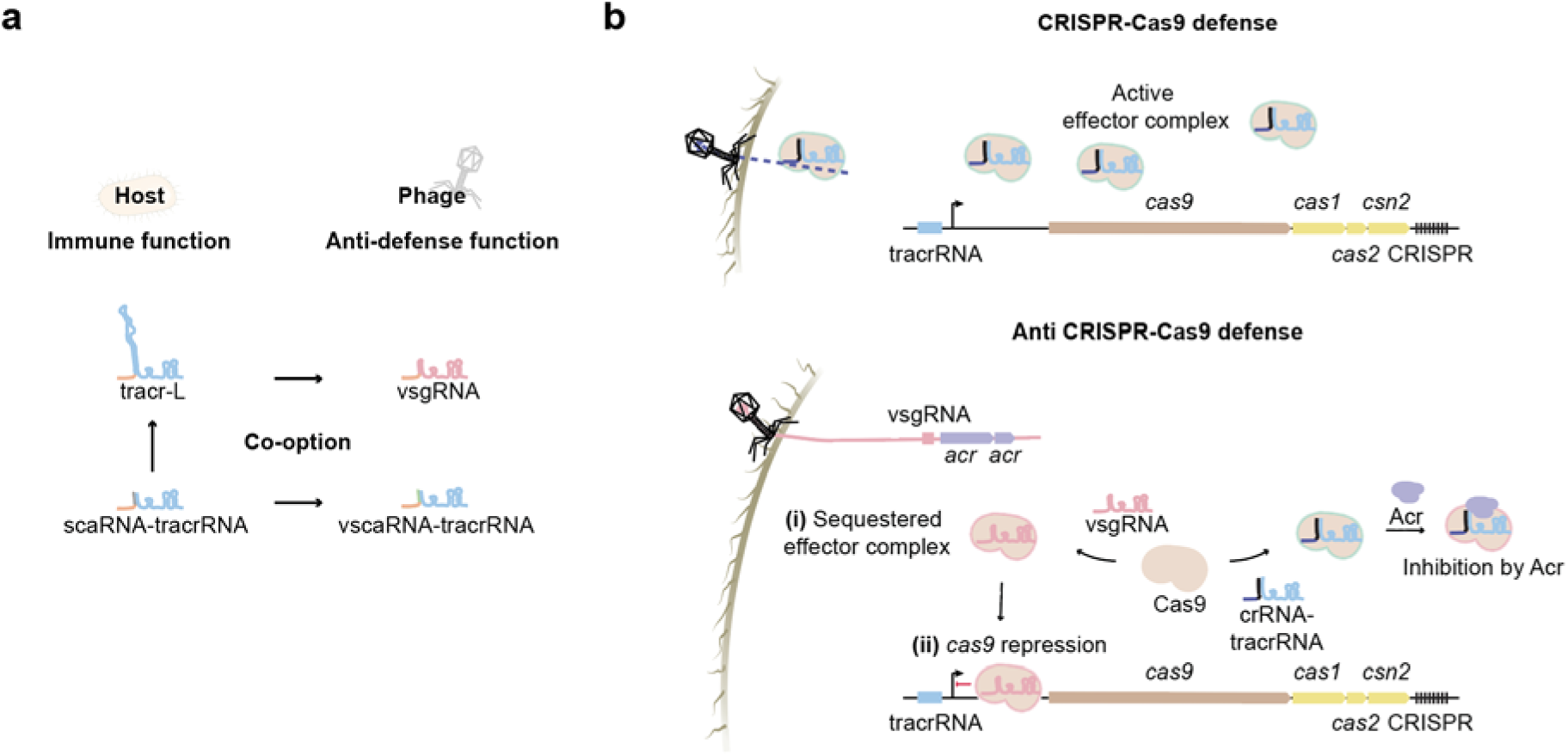
Model for phage-mediated Cas9 inhibition via the co-option of Cas9-regulating RNAs. a, Phages acquire host-derived tracr-L and scaRNA as anti-defense non-coding RNAs. **b,** These co-opted ncRNAs neutralize Cas9 via two mechanisms: (**i**) sequestration, in which the ncRNAs competitively bind Cas9 to preclude the assembly of interference-competent tracrRNA-crRNA complexes; and (**ii**) transcriptional repression, in which Cas9 is guided to its own promoter, leading to *cas9* operon silencing and the prevention of downstream gene expression.

### Modified vsgRNAs enable gene editing in human cells

Apart from subverting Cas9 immunity in bacteria, vsgRNAs could directly offer naturally occurring and compact guide RNAs for CRISPR technologies. To this end, we explored the extent to which vsgRNAs could enact Cas9-mediated gene editing in mammalian cells. We first evaluated indel formation through targeted DNA cleavage with a delivered Cas9-Hifi ribonucleoprotein complex^39^. To enable DNA cleavage by the vsgRNA, we extended the guide portion to the standard 20 nt, with the resulting RNA still 12 nts shorter than a standard sgRNA (**Fig. S15a**). Under this setup, the vsgRNA enabled indel formation (10% − 93%) across all five tested sites in four cell lines (**Fig. S15b**). While the editing frequency was lower than that of an equivalent sgRNA, this gap was overcome and even exceeded by introducing stabilizing mutations to the vsgRNA lower stem as well as RNase-resistant chemical modifications^40^ (GOLD vsgRNA, **Figs. S15a-b**). All guide RNAs resulted in negligible editing (≤ 0.1%) at the top two predicted off-targets for each site^41^ (**Fig. S15b**). The modified vsgRNA could also enact adenine base editing via RNP delivery^42^, with similar trends to indel formation (**Fig. S15c**). Finally, the vsgRNA could enact indel formation when expressed from a U6 promoter at two difficult-edit sites in an induced pluripotent cell line, with the stem-stabilizing mutations improving the editing efficiency (**Fig. S15d**). In total, these results show that vsgRNAs can be readily exploited for genome editing.

## DISCUSSION

Here we reveal that phages encode sgRNA-like molecules we dubbed vsgRNAs that subvert Cas9 immunity. vsgRNAs can act at two levels of inhibition: sequestering Cas9 holoenzymes that can no longer participate in immune defense, and directing Cas9 to downregulate its own transcription (**Fig. 5**). The latter level involves a short guide region that directs Cas9 to bind, but not cleave, its own promoter to block transcription of *cas9* and any downstream genes in the operon. In our experiments, CRISPR immunity was inhibited principally through Cas9 sequestration (**Fig. 2b,f**), although even modest repression of *cas* expression can greatly impair immune defense^43,44^. As *cas9* and the downstream genes are involved in spacer acquisition^45,46^, silencing may also impinge on adaptive immunity. Beyond vsgRNAs, we discovered a distinct class of inhibitors we termed vscaRNAs that should act similarly to the vsgRNAs while also sequestering endogenous tracrRNA needed to form the dual-guide RNA bound by Cas9. Bioinformatic evidence indicates that both inhibitors were co-opted from Cas9-regulated RNAs associated with diverse CRISPR-Cas9 systems based on shared guide sequences and tracrRNA tails (**Fig. 5a**). These inhibitors parallel the recent discoveries of phage-encoded Racrs that mimic CRISPR repeats to subvert immunity through Cas sequestration alone^19,24^ or through directed silencing of CRISPR-Cas expression^47^. The vsgRNAs also form a broad set of bacterial defenses hijacked by phages for their own purposes^18^, such as Cas proteins converted into Acrs^48^ or non-coding RNAs that inhibit an anti-phage toxin-antoxin system^49^. While our analyses identified 170 vsgRNAs associated with type II-A and II-C CRISPR-Cas systems, many more likely remain undiscovered due in part to our stringent requirement for a tracrRNA within the same genome.

vsgRNAs were consistently encoded with Acrs, comprising expansive anti-CRISPR islands^23^. While Acrs are known that would be expected to interfere with both inhibitory mechanisms of vsgRNAs^33,50^, the co-encoded Acrs we identified in our bioinformatics search should allow for both Cas9 sequestration and, in most cases, also auto-repression. For example, CasPRs block transcription by binding within *cas* coding regions that do not overlap with vsgRNA target sites within the promoter region^31^, while AcrIIA18 truncates crRNA guides to 15 nts that still exceeds the length of the vsgRNA guide^33^. Most interestingly, AcrIIA25.1 blocks DNA binding by sgRNA-loaded Cas9^21^ but has no effect on the vsgRNA-loCas9 complex due to mutations within the loop regions of the vsgRNA, paralleling Racrs regulating CRISPR-Cas expression that escape inhibition by AcrIF24^47^. AcrIIA27 represents one of the few examples of co-encoded Acrs that could inhibit DNA binding by vsgRNAs^34^, although our work suggests that Cas9-vsgRNA binding alone primarily accounts for immune inhibition. Anti-CRISPR islands thus may not be random collections of inhibitors but instead provide complementary mechanisms to inhibit immune defense.

The evolution of vsgRNAs parallels the engineering of sgRNAs and other RNA-guided technologies. sgRNAs were invented by fusing the processed crRNA and tracrRNA without modifications to the retained sequence, with the goal of producing a single RNA and jettisoning the need for RNase III. Later work showed that mutations or chemical modifications to the sgRNA could improve gene editing efficiencies^40,51^, while truncating the guide sequence or introducing PAM-distal mismatches enabled DNA binding with Cas9 otherwise capable of DNA cutting^38,52,53^. Similarly, the vsgRNA likely emerged when phages assimilated tracr-L followed by deleting the elongated upper stem and mutating the lower stem to prevent pairing with the pre-crRNA^6^, and mutating the tracrRNA tail to prevent some Acrs from inhibiting DNA binding by the Cas9-vsgRNA complex (**Figs. 1c,f, 3c and 5a**). The co-opted tracr-L then likely underwent optimization to suppress immunity by condensing the RNA while retaining Cas9 binding and promoter targeting and also preventing inhibition by Acrs that block DNA binding. Beyond sgRNAs and vsgRNAs, other parallel examples reflected engineering preceding discovery. For instance, Cas nucleases were engineered for gene repression or activation by introducing catalytically inactivating mutations and fusing transcriptional activating or repressing domains^54^, while recent work revealed natural counterparts exapted from CRISPR-Cas systems and transposons that are directed by encoded guide RNAs to activate or repress gene expression^55,56^. Separately, riboswitches were originally engineered by coupling *in vitro*-selected aptamers to ribozymes followed by other functional domains to enable ligand-dependent control of cellular processes^57,58^, while natural riboswitches were subsequently discovered that provide feedback regulation of numerous core metabolic processes and are considered surviving fossils from ancient organisms^59^. These examples underscore how evolution can parallel human design and suggest that other purely engineered biomolecular systems may have yet-to-be-discovered natural counterparts.

Investigating the prevalence and diversity of vsgRNAs led us to discover Cas9-regulated RNAs most similar to scaRNAs (**Fig. 5a**). While the scaRNA was previously found in only a single bacterium and shown to regulate host virulence genes^16,17,38^, we found similar RNAs that instead regulate Cas9 expression. Like tracr-L, the Cas9-regulating scaRNAs would suppress Cas9 expression that could be reversed when needed, such as the presence of phage-encoded Acrs that suppress DNA cleavage and binding or in a small sub-population of cells that overexpresses Cas9 through mutation of the scaRNA target site^6,20^. The discovery of cas9-regulating scaRNAs in different locations within the CRISPR-Cas system (**Fig. S11**) suggests that tracr-L represents a scaRNA located upstream of the tracrRNA. Despite some differences between tracr-L and Cas9-regulating scaRNAs, both modes of regulation would be convenient for phages to hijack as we observed with vsgRNAs and vscaRNAs (**Fig. 5a**), potentially explaining why these modes of RNA-based negative feedback regulation of Cas9 are not more prevalent (**Fig. 4a**). Finally, the two classes of Cas9-regulating RNAs add to the set of known regulatory RNAs expressed from within CRISPR-Cas systems that suppress the transcription of co-encoded toxins or complementary immune systems^60–63^, underscoring how evolution has taken advantage of Cas effectors to enact gene regulation separate from immune defense.

## METHODS

### Primers, plasmids, and growth conditions

The strains, DNA oligonucleotides, plasmids, RNAs, and gBlocks used in this study are listed in **Tables S7-S11**, respectively. In general, genes of interest were amplified from *Streptococcus oralis* subsp. *tigurinus* 859 or *Lactobacillus hominis* DSM 23910 genomic DNA using Q5 Hot Start High-Fidelity DNA Polymerase (NEB), or ordered as gBlocks (IDT), and then assembled into plasmids using NEBuilder HiFi DNA Assembly Master Mix (NEB). Mutations were introduced by PCR using Q5 Polymerase. All constructs were verified by DNA sequencing.

*E. coli* was grown at 37°C in Lysogeny Broth (LB) medium with shaking at 220 rpm or on LB agar plates (LB broth, 18 g/l agar), except for strain KL740 cI857+, which was cultured at 29°C. *E. coli* strains TOP10, DH5α, or KL740 cI857+ were used for plasmid cloning. Antibiotics were added where appropriate at final concentrations of 100 μg/ml for ampicillin, 34 or 50 μg/ml for chloramphenicol, and 50 μg/ml for kanamycin for *E. coli*. *Streptococcus oralis* subsp. *tigurinus* strain 859 (*S. tigurinus* 859) was grown in Tryptic Soy Broth (TSB) at 37°C with no shaking. *Lactobacillus hominis* DSM 23910 was grown in De Man, Rogosa, and Sharpe medium (MRS) at 37°C.

### Northern blotting analysis, 5**′** RACE, and RNA-seq

For total RNA extraction from *Streptococcus tigurinus* 859, overnight culture of *Streptococcus tigurinus* 859 was grown in TSB from a single colony at 37°C without agitation. Then the culture was diluted to OD_600_ 0.05 and grown in TSB to OD_600_ 0.4-0.6, and 5 ml of culture was mixed with 1 ml of stop solution (95% v/v EtOH + 5% v/v phenol), harvested by centrifugation at 5,000 x g for 10 min at 4°C. Another aliquot of 5 ml was harvested the next day at OD_600_ 1.3-1.4 the same way. Cell pellet was resuspended in 600 µl of 5 mg/ml lysozyme in 10 mM Tris-EDTA pH 8.0; 60 µl 10% w/v SDS was added, the mix was incubated at 64°C for 1 min. To the cell lysate, 66 µl 3M NaOAc, pH 5.2 and 750 µl phenol were added, the mix was incubated at 64°C for 6 min. For the phase separation, the samples were centrifuged at 15,000 x g for 15 min at 4°C.

The aqueous layer was transferred into new tubes and mixed with 750 µl of chloroform. After another centrifugation at 15,000 x g for 15 min at 4°C, the aqueous layer was mixed with 1.4 ml (30:1) EtOH:NaAcetate (from a 3 M stock at pH 6.5). After the samples were placed at −20°C for RNA precipitation overnight, they were centrifuged at 15,000 x g for 30 min at 4°C. The supernatant was removed, and the RNA pellet was washed with 70% EtOH and air dried. RNA pellet was resuspended in water and cleaned with RNA Clean and Concentrator columns-25 (Zymo Research). For the 5′ RACE and RNA-seq, RNA was treated with DNase I (Zymo Research).

A total of 500 ng of DNase treated total RNA extracted from *Streptococcus tigurinus* 859 was subjected to 5′ RACE using the Template Switching RT Enzyme Mix (New England Biolabs). Reverse transcription was performed with RT primer St859_vsgRNA_NB_RT. To create a cDNA library, PCR was performed using the Q5 Hot Start High-Fidelity 2X Master Mix (New England Biolabs) with the PCR primer St859_vsgRNA_5RACE_PCR and a TSO-specific primer TSO-specific_primer. The cDNA libraries were purified using Zymo DNA Clean and Concentrator columns-5 (Zymo Research) cloned into the supplied linearized vector pMiniT 2.0 using NEB PCR Cloning Kit (New England Biolabs). In total, 16 random colonies were picked for colony PCR with the primers 5RACE_cPCR_Fw and 5RACE_cPCR_Rv to determine sequence of the plasmid insert, purified using Zymo DNA Clean and Concentrator columns-5 (Zymo Research) and submitted for Sanger sequencing.

For the Northern blotting analysis, 10 μg of total RNA extracted from *Streptococcus tigurinus* 859 was mixed with 2x RNA loading dye (0.025% w/v bromophenol blue, 0.025% (w/v) SDS, 0.025% w/v xylene cyanol, 18 mM EDTA (pH 8.0), 93.64% v/v formamide), denatured at 95°C for 5 min, quickly placed on ice and loaded on 8% PAAG-7M urea gel. RNA was transferred from a gel onto Hybond-XL membranes (Amersham Hybond-XL, GE Healthcare) using an Electroblotter with an applied voltage of 50 V for 1 h at 4°C (Tank-Elektroblotter Web M, PerfectBlue), crosslinked twice with 120 mJ/cm2 UV-light (UV-lamp T8C; 254 nm, 8 W), hybridized overnight in Roti-Hybri-Quick buffer at 42°C with ^32^P-5′ end-labeled oligodeoxyribonucleotides. St859_vsgRNA_NB_RT was used for the vsgRNA detection. After washing first with 5x and then with 1x SSC buffer, the membrane was exposed to a storage phosphor screen and visualized using an Amersham Typhoon imaging system. The assay was performed in three biological replicates.

For total RNA extraction from *Lactobacillus hominis* DSM 23910, the culture was growing from a single colony in MRS medium (ROTH) in a candle jar for 4 days, then the culture was harvested. RNA extraction and DNase I treatment steps are the same as for *Streptococcus tigurinus* 859 described earlier.

cDNA libraries for RNA-seq were prepared from DNase treated total RNA from *Streptococcus tigurinus* 859 using Illumina Stranded Total RNA Prep with Ligation and Ribo-Zero Plus Microbiome reagents. DNA libraries for sRNA-seq were prepared from DNase-treated total RNA from *Streptococcus tigurinus* 859 or *Lactobacillus hominis* DSM 23910. To ensure efficient primer ligation to the 3′-ends for sRNA-seq, the RNA was treated with PNK (NEB) for 30 min at 37°C, then the RNA was purified with Direct-zol™ RNA MiniPrep using TRI Reagent™ (Zymo Research). Then, the libraries were prepared using Takara SMART-seq sRNA; size selection was performed according to the Takara protocol to enrich for inserts shorter than 150 bp. The libraries were sequenced on an Element Biosciences AVITI sequencer, using paired-end 75 bp reads, with a total yield of up to ∼100 million reads. FASTQ files were processed using the Galaxy Europe platform (https://usegalaxy.eu). The reads were mapped with bowtie2^64^ with the very sensitive end-to-end preset to the genome with the vsgRNA (LNVH01000004.1) or *cas9* (LNVH01000012.1) loci in *S. tigurinus* or scaRNA/*cas9* locus (CAKE01000018.1) in *L. hominis*. The mapped reads were converted into WIGGLE files^65^ and visualized with IGV_2.19.7.

### Nucleic acid labeling

For the 5′ end labeling of RNA/DNA with ^32^P used for the Northern blotting/in-line probing, synthetic oligoribonucleotides/oligodeoxyribonucleotides (IDT) were treated with T4 PNK (New England Biolabs) in the presence of [gamma-^32^P]ATP (Hartmann Analytic). The labeled DNA was purified on Sephadex G25 column, eluted in water. The labeled RNA was mixed with 2x RNA loading dye (0.025% w/v bromophenol blue, 0.025% w/v SDS, 0.025% w/v xylene cyanol, 18 mM EDTA (pH 8.0), 93.64% v/v formamide), separated on a 8% polyacrylamide gel (PAAG)-8M urea; the band with the RNA was excised from the gel, and the RNA was extracted from it using ZR sRNA PAGE Recovery Kit (Zymo Research).

For the 3′ end labeling of RNA with Cy5 used for EMSA, RNAs were treated with T4 RNA ligase (Thermo Fisher Scientific) in the presence of pCp-Cy5 (Jena Bioscience). The labeled RNA was purified with RNA Clean and Concentrator columns-5 (Zymo Research).

For in vitro binding and cleavage assays, the 5′ ends of the oligodeoxyribonucleotides were labelled with Cy5 using 5′ EndTag™ DNA/RNA Labeling Kit (Vector Laboratories) and Cy®5 Maleimide Mono-Reactive Dye (Cytiva). The labeled DNA was purified with DNA Clean and Concentrator columns-5 (Zymo Research).

### In-line probing

For in-line probing sample, ∼0.2 pmol ^32^P-5′ end-labeled vsgRNA (St859_vsgRNA in the **Table S10**) was incubated for 40h at room temperature in an in-line probing buffer (50 mM Tris-HCl, pH 8.3, 20 mM MgCl_2_, and 100 mM KCl). For alkaline ladders (OH), ∼0.2 pmol ^32^P-5′ end-labeled RNA was denatured for 5 min at 95°C in alkaline hydrolysis buffer (Ambion). For RNase T1 ladders, 0.2 pmol ^32^P-5′ end-labeled RNA and 10 μg of yeast RNA (Ambion) was incubated in 1x sequencing buffer (Ambion) for 1 min at 95°C, transferred on ice and further incubated with 0.1 U RNase T1 (Ambion) for 5 min at 37°C. All reactions were prepared in a 10-μl volume and were stopped by adding 10 μl colorless gel-loading solution (10 M urea, 1.5 mM EDTA, pH 8.0) on ice. The reactions were analyzed on 15% PAAG-8M urea. The gels were dried, exposed to a storage phosphor screen, and analyzed using an Amersham Typhoon imaging system. The assay is performed in two replicates.

### PAM prediction

Primers were designed to flank the known CRISPR array, followed by PCR amplification of genomic DNA. The purified PCR product was then subjected to Sanger sequencing to bridge the gaps between disconnected contigs. Finally, 30 spacer sequences in the CRISPR array adjacent to *cas9* gene were retrieved, and searched against the NCBI nr database using BLASTN (*E*-value cutoff 0.05, minimum 90% identity, word size 11) to identify putative protospacers. Up to 12 nts of downstream flanking sequences were further retrieved for each putative protospacer to capture potential PAM sequences. Protospacers with flanking sequences were then aligned to each other using MUSCLE^66^, and visualized using WebLogo (https://weblogo.threeplusone.com/create.cgi) to analyze PAM conservation patterns.

### Phage plaque assays

Phage T7 was propagated on *E. coli* MG1655 ΔRM by picking a single phage plaque into a liquid mid-log culture. The culture was then grown for 8 h at 37°C, after which the supernatant was harvested by centrifugation at 5,000 rpm for 10 min and subsequently filtered through a 0.22 µm filter to remove bacterial contamination.

Phage plaque assays were performed via the double agar overlay method. *E. coli* MG1655 ΔRM host strains carrying the corresponding plasmids were grown overnight in LB medium supplemented with appropriate antibiotics and inducers. Typically, for pBAD-Cas9 plasmids, 50 µg/mL chloramphenicol and 0.2% w/v L-arabinose were added. For gRNA plasmids, 100 µg/ml ampicillin was added. For vsgRNA plasmids, 50 µg/ml kanamycin and 1 mM isopropyl-ß-d-thiogalactopyranoside (IPTG) were added. Then, 100 µl of overnight culture was mixed with 5 ml melted 0.75% w/v top agar containing the appropriate supplements and then poured onto a pre-poured 1.5% w/v LB agar plate with the same supplements and left to solidify. Phages were 10-fold serial diluted in phage buffer (10 mM Tris-HCl, pH 7.4, 10 mM MgSO_4_, 0.01% w/v gelatine), and 5 µl of each dilution were spotted onto the top agar overlay. Plates were imaged and plaque forming units (PFUs) were counted after overnight incubation at room temperature.

### Fluorescent reporter assays

Bacterial host strains with corresponding plasmids from a single colony were grown overnight in LB medium supplemented with appropriate supplements. The overnight cultures were then diluted in fresh LB medium plus appropriate supplements to a final OD_600_ of 0.02. Then, 200 µl of the cultures were transferred into a 96-well plate and kept at 37°C with continuous shaking in the Varioskan LUX multimode microplate reader (Thermo Scientific), and the fluorescence and OD_600_ were measured every 10 min for a total duration of 20 h. For each experimental group, at least three independent colonies were randomly selected and cultured.

### In vitro transcription

For the in vitro RNA binding or DNA binding/cleavage assays, sgRNA was synthesized using HiScribe T7 High Yield RNA Synthesis Kit (New England Biolabs). The T7 transcription template was ordered as a gBlock St859_sgRNA-T7-gB (IDT). The transcription was treated with DNase I (Zymo Research) and cleaned with RNA Clean and Concentrator columns-25 (Zymo Research).

### In vitro RNA binding assay

For the electrophoretic mobility shift assay (EMSA), 2 nM Cy5-labeled RNA was incubated with various amounts of SpCas9 (New England Biolabs) from 1 nM to 40 nM for 15 min at 37°C in a 1x NEBuffer™ r3.1 (New England Biolabs) in the presence of 125 ng/μL of yeast RNA (#AM2283, Ambion) to enhance specific RNA-protein interaction. The samples (10 μL) were separated on 2% agarose-TBE gels at 4°C. Gels were visualized using an Amersham Typhoon imaging system. The assay is performed in three replicates. The Kd was calculated using GraphPad Prism 11 (non-linear fit, one site, specific binding).

### In vitro DNA binding and cleavage assay

dsDNAs containing the vsgRNA target site with PAM or mutated PAM were created by annealing the oligodeoxyribonucleotides St859_promoter_Fw/St859_promoter_Rv and St859_mutpromoter_Fw/St859_mutpromoter_Rv. The oligodeoxyribonucleotides St859_promoter_Fw and St859_mutpromoter_Fw were labelled with Cy5 before annealing. Annealing was performed by mixing the oligodeoxyribonucleotides in equal amounts at 65°C for 13 min, then 10 min at room temperature.

The samples for the assay were prepared by incubating the 125 nM Cy5-labelled dsDNA with 1 μM vsgRNA or sgRNA and 40 nM SpCas9 (New England Biolabs) for 15 min at 37°C in a 1x NEBuffer™ r3.1 (New England Biolabs) in the presence of 125 ng/μL of yeast RNA (#AM2283, Ambion) to enhance specific protein-nucleic acid interaction. The samples (10 μL) were separated on 2% agarose-TBE gels at 4°C to monitor binding of the RNP to the DNA or were mixed with 10 μl of colorless gel-loading solution (10 M urea, 1.5 mM EDTA, pH 8.0) and separated on 10% polyacrylamide gel with 7 M urea to monitor cleavage of the DNA. Gels were visualized using an Amersham Typhoon imaging system. The assay is performed in three replicates.

### Cell-free transcription-translation assays

Plasmids encoding either the Cas9 nuclease, a gRNA, an Acr, or GFP were used to assess the targeting activity in a cell-free transcription-translation assay. SpCas9 plasmid (CBS-119) was previously constructed^67^. LhCas9 plasmid (CBB-0090) was constructed with Gibson Assembly; the gene of *cas9* was amplified from the *L. hominis* DSM 23910 gDNA. Plasmids with Acrs (CBB-0091, CBB-0092, CBB-0093) were constructed with Gibson Assembly; the Acr genes were amplified from *S. tigurinus* 859 gDNA. To construct targeted plasmids, target sequences and PAM were inserted directly downstream of the constitutive OR2-OR1 promoter (CBB-0094, CBB-0097) or downstream of the deGFP CDS (CBB-0095) using Q5 site-directed mutagenesis. gRNAs were constructed either with Gibson Assembly or with Q5 site-directed mutagenesis. The plasmids were isolated from *E. coli* using ZymoPURE II Plasmid Midiprep Kit (Zymo Research) and purified with DNA Clean and Concentrator columns-25 (Zymo Research).

To assess binding of vsgRNA to the promoter, plasmids encoding SpCas9 (CBS-119), a gRNA (vsgRNA - CBB-0098, tracr-L - CBB-0099, sgRNA - CBB-0100, or NT sgRNA - CBB-0101), and a target-GFP (CBB-0094 or CBB-0096) in final concentrations of 3 nM, 0.5 nM, and 0.5 nM, respectively, were mixed in 3 μL of water, then 9 μL of myTXTL master mix (Daicel Arbor Biosciences) was added. 5 μL of the sample were then incubated at 29°C for 16 h in a plate reader (Agilent BioTek Synergy H1) and fluorescence was measured after 16 hours (excitation, emission: 485 nm, 528 nm). The values of the GFP fluorescence (AU) were scaled by a factor of 1,000 for visualization and plotted with GraphPad Prism 11. The assay was performed in four replicates.

To assess repression of targeting with Acrs, Acrs were first pre-expressed in TXTL. For that, 3 μL of a plasmid encoding an Acr (AcrIIA25.1 - CBB-0091, AcrIIA24 - CBB-0092, or AcrIIA3 - CBB-0093) was mixed with 9 μL of myTXTL master mix to a final concentration of *acr*-encoding plasmid of 18 nM, and then incubated at 29°C for 16 h. Plasmids encoding SpCas9, a gRNA, and a target-GFP (CBB-0094 or CBB-0095) in final concentrations of 0.5 nM each, were mixed in 2 μL of water, then 1 μL of the pre-expressed Acr and 9 μL of myTXTL master mix (Daicel Arbor Biosciences) were added. 5 μL of the sample were then incubated at 29°C for 16 h in a plate reader (Agilent BioTek Synergy H1) and fluorescence was measured after 16 hours (excitation, emission: 485 nm, 528 nm). The assay was performed in four replicates. Similarly, mutations of sgRNA in the stem-loops (SL1 - CBB-0102, SL2 - CBB-0103, SL3 - CBB-0104, SL4 - CBB-0105) or guide (CBB-0106) were tested. The values of the GFP fluorescence (AU) were scaled by a factor of 1,000 for visualization and plotted with GraphPad Prism 11. The assay was also performed in four replicates.

To assess binding of scaRNA-tracrRNA to the promoter, the scaRNA, tracrRNA, NT crRNA, and T sgRNA were in vitro transcribed using HiScribe® T7 Quick High Yield RNA Synthesis Kit (New England Biolabs). The sequences of the RNAs are in **Table S10**. scaRNA-tracrRNA (14 μM each), NT crRNA-tracrRNA (14 μM each), T sgRNA were denatured at 72°C for 3 min, then annealed by incubating at room temperature for 20 min. To form an RNP with LhCas9, LhCas9 was pre-expressed in TXTL. For that, 1 μL of a plasmid encoding LhCas9 (36 nM) (CBB-0090) was mixed with 9 μL of myTXTL master mix and incubated at 29°C for 4 hours. After 1.5 μL of the RNAs (14 μM) were added to the TXTL with pre-expressed LhCas9, the tubes were incubated at room temperature for 15 min to allow the RNPs to form. After that, the plasmid with the target-GFP (CBB-0097 or CBB-0108) was added in final concentration of 0.5 nM. 5 μL of the sample were then incubated at 29°C for 16 h in a plate reader (Agilent BioTek Synergy H1) and fluorescence was measured after 16 h (excitation, emission: 485 nm, 528 nm). The values of the GFP fluorescence (AU) were scaled by a factor of 1,000 for visualization and plotted with GraphPad Prism 11. The assay was performed in four replicates.

To assess binding of vscaRNA-tracrRNA to the promoter, the vscaRNA with 11 nt or 20 nt guide, tracrRNA, NT crRNA, and crRNA with 20 nt guide were in vitro transcribed using HiScribe® T7 Quick High Yield RNA Synthesis Kit (New England Biolabs). The sequences of the RNAs are in **Table S10**. vsca(cr)RNA-tracrRNA (14 μM each) duplexes were denatured at 72°C for 3 min, then annealed by incubating at room temperature for 20 min. To form an RNP with LhCas9, LhCas9 was preexpressed in TXTL. For that, 1 μL of a plasmid encoding LhCas9 (36 nM) (CBB-0090) was mixed with 9 μL of myTXTL master mix and incubated at 29°C for 4 hours. After 1.5 μL of the RNA duplexes (14 μM) were added to the TXTL with pre-expressed

LhCas9, the tubes were incubated at room temperature for 15 min to allow the RNPs to form. After that, the plasmid with the target-GFP (CBB-0168 or CBB-0169) was added in final concentration of 0.5 nM. 5 μL of the sample were then incubated at 29°C for 16 h in a plate reader (Agilent BioTek Synergy H1) and fluorescence was measured after 16 h (excitation, emission: 485 nm, 528 nm). The values of the GFP fluorescence (AU) were scaled by a factor of 1,000 for visualization and plotted with GraphPad Prism 11. The assay was performed in four replicates.

### Identifying natural sgRNA based on the experimentally validated and predicted examples

A covariance model was built from each of 79 previously characterized sgRNA and one predicted sgRNA using the Infernal software (version 1.1.3) with the command cmbuild. These 80 sgRNAs represent the high diversity of all previously known Cas9 families^30^. With these models, a non-redundant set of predicted sgRNAs^68^ was searched using cmsearch^69^ with a stringent E-value threshold of 10^-10^. If no hit was found in the predicted set of sgRNAs, the covariance model with the validated sgRNA was used for the further steps. Otherwise, out of the obtained alignments, new diversified covariance models were built with cmbuild.

The bacterial genomes from the RefSeq database (version 223)^70^ were downloaded from the FTP server (https://ftp.ncbi.nlm.nih.gov/genomes/refseq). The downloaded genomes were searched with the diversified covariance models with the cmsearch^69^ program from the Infernal package (Version 1.1.3) using an E-value cutoff of 0.1. Moreover, the 80 covariance models were also used to search the IMG/VR database (Version 7.1)^71^ with the command cmsearch and an E-value cut off of 0.001.

### Identifying vsgRNAs in the RefSeq and IMG/VR databases

The results of the homology search against the RefSeq database were filtered to include only genomes containing more than one hit of the covariance models. If one natural sgRNA showed a maximum of one mismatch within the first seven base pairs of the repeat:anti-repeat stem-loop—as predicted by covariance models—and was located more than 10 kb away from the *cas9* gene, while the other covariance model hit was located within 10 kb of *cas9*, then the distal natural sgRNA was defined as a vsgRNA.

Similarly, natural sgRNA candidates identified in the IMG/VR database were screened for the same stem-loop structural constraints. The viral contigs of these potential vsgRNA were checked for the presence of *cas9*. First, all proteins of each contig were annotated with Prokka_v (version 1.12-viral)^72,73^ and then analyzed with the command hmmsearch from the HMMER software package (Version 3.4)^72^ and an alignment of 11 Cas9 proteins downloaded from the NCBI conserved domain database^74^ (https://www.ncbi.nlm.nih.gov/Structure/cdd/TIGR01865, February 2024).

Every natural sgRNA in the IMG/VR database without *cas9* in the same viral contig and which was smaller than 120 nts was defined as vsgRNA.

### Predicting tracr-L with *cas9* promoter targets

The results of the homology search against the RefSeq database were filtered into two categories: normal and long tracrRNAs. TracrRNA hits with a maximum of one erroneous base pair (all base pairs that are not A-U, G-C or G-U) within the initial 11 bp of the repeat:anti-repeat stem-loop, as predicted by the covariance models, were identified as potential tracr-L. The remaining sequences were assumed to be short tracrRNAs. Following the initial filtering stage, the RefSeq annotation of the genome in which the potential tracr-L was identified was examined to ascertain the presence of a *cas9* gene within 10 kb of the tracrRNA based on the annotation of the NCBI nuccore database^75^.

If both conditions were met, the 12-nt spacer sequence of each potential tracr-L was extracted, and a perfectly matching target site of at least 11 nts between the potential tracr-L and the start codon of *cas9* was sought. If the potential tracr-L was downstream of *cas9*, 500 nts upstream of *cas9* start codon were searched instead. Only if all three conditions were met was the tracrRNA considered to be a tracr-L (**Table S4**).

### Identifying scaRNA targeting *cas9*

For each tracrRNA predicted with the previously described pipeline that had an E-value of below 0.001, all annotated genes with 10 kb on both sides were downloaded from the NCBI nuccore database^75^. If a CRISPR array was present within the same CRISPR-Cas locus, the intergenic regions of the CRISPR-Cas locus and the subsequent genes were extracted and downloaded. The strand of the CRISPR array was predicted by calculating the minimum free energy of the interaction between the repeat sequence (or the repeat sequence of the opposite strand) and the tracrRNA sequence with the IntaRNA (Version 3.2.0) software^76^. The strand with the lowest minimum free energy between the tracrRNA and the repeat was assumed to be the correct one^77,78^.

Once the correct repeat sequence had been identified, the intergenic regions were searched for repeat-like sequences using the BLASTN tool (version 2.12.0+, with the default parameters except for -wordsize 5 and -evalue 0.01)^79^. Hits were filtered based on their length (minimum 20 nts), e-value (less than 0.001), and sequence identity (greater than 80%), with the additional criterion that the hits should include the first nucleotide of the repeat.

Subsequently, a potential target site was sought within the CRISPR-Cas locus by BLASTing (using the same parameters except -evalue 100) the 15 nts upstream of the repeat-like element against the intergenic regions. All hits with a length of below 10 nts, a sequence identity of less than 100%, or a missing 3’ end of the spacer were excluded from further analysis. The remaining hits were then subjected to manual curation to ascertain whether they exhibited a matching PAM sequence (based on the closest characterized Cas9 from prior work^30^) and whether they targeted the predicted transcription starting site^80^ (40 nts upstream and 20 nts downstream). If both conditions were met, the repeat-like element was assumed to be a scaRNA that targets *cas9*.

### Search for viral scaRNAs

To identify viral scaRNA, six representative, non-redundant scaRNA sequences were queried against the RefSeq database using the BLASTN web tool (NCBI; accessed March 18, 2026). Searches utilized the “Somewhat similar sequences” algorithm (E-value threshold: 1000) with a truncated query comprising the 11-nt spacer and the first 7 nt of the scaRNA repeat. Note that the entire scaRNA repeat was not used to avoid BLAST retrieving the repeats from the CRISPR array. Genomes harboring at least two hits were used for further analysis; a viral scaRNA was defined as a hit located >10 kb from a *cas9* locus in a genome where a second “proximal” hit resided within 10 kb of *cas9*, mirroring the spatial filtering criteria used for vsgRNAs.

Additionally, the IMG/VR database (Version 7.1) was queried using standalone BLASTN (v2.12.0+; parameters: -wordsize 5, -evalue 0.05). In contrast to the RefSeq search, these queries utilized the full-length repeat-like elements. Candidates were classified as viral scaRNAs if they exhibited an E-value <0.05 and 100% sequence identity across the first 10 nt of the spacer region.

### Analysis of the neighborhood of vsgRNAs and vscaRNAs

For all vsgRNA and vscaRNA found in the RefSeq database, all proteins 2kb up- and downstream from the vsgRNA were downloaded from the NCBI nuccore database^75^. For the vsgRNAs and vscaRNA found in the IMG/VR database, all proteins 2kb up- and downstream of the vsgRNAs were extracted from the above-annotated viral contigs.

As a next step, all anti-CRISPR proteins within the extracted proteins were annotated with hmmscan (standard parameters) from the HMMER software package (Version 3.4)^72^ and the anti-defense finder database^81^ (https://github.com/mdmparis/defense-finder-models/tree/master/profiles, downloaded 06.11.2025). A protein was identified as an anti-CRISPR protein if the E-value assigned by hmmscan was below 10^-5^.

Moreover, we utilized a set of previously predicted anti-CRISPR proteins^82^ to identify potential anti-CRISPR proteins near the vsgRNA. For that, we took all predicted anti-CRISPR proteins from prior work^82^ with a score higher than 0.5 and used the blastp tool (standard parameters except -evalue 0.01) from the BLAST+ software package (version 2.12.0+) to search homologs in the proteins next to the vsgRNAs. All proteins with a blastp hit which had an E-value below 10^-5^ were defined as predicted anti-CRISPR proteins. The remaining proteins were defined as non-anti-CRISPR proteins. The same blastp parameters were used to identify homologs to the newly identified Gut_Acr^83^ and CasPR^31^.

To determine if the candidate vsgRNAs and vscaRNAs identified in RefSeq were localized within integrated prophages, the corresponding bacterial genomes and contigs were retrieved from the NCBI Nucleotide (nuccore) database. These sequences were subsequently analyzed using GeNomad (v1.8.0, default parameter with end-to-end)^84^ to identify and classify proviral regions harboring the ncRNA candidates.

### Guide comparison between vsgRNA and the tracr-L

To assess potential targeting homology, the 10-nucleotide (nt) sequences immediately upstream of all identified vsgRNAs were extracted. These flanking regions were queried against the first 10 nt of the guide sequences from approximately 1,800 tracr-L candidates (identified in RefSeq) using standalone BLASTN (v2.12.0+). To account for the short query length, the alignment parameters were optimized for high-sensitivity searches (-word_size 5, -evalue 100, - gapopen 3, -gapextend 5, -penalty -5, -reward 4). For sequences failing to return a significant

BLAST hit, a secondary pairwise alignment was performed to manually quantify nucleotide-level mismatches at each position.

### Visualizing sequence and secondary structure of RNAs

Schematic RNA secondary structures were illustrated using https://rna2drawer.app/ and RNA secondary structure with nucleotide resolution were visualized with R2R^85^.

### Mammalian cell culture

Mammalian cells were grown at 37°C in a humidified incubator with 5% CO_2_. We used 409-B2 human induced pluripotent stem cells (Riken BioResource Center, HPS0076, GMO permit AZ 54-8452/26) with or without inducible Cas9^86^; HEK293 cells (ECACC, 85120602) grown in DMEM/F-12 (Gibco, 31330-038) supplemented with 10% fetal bovine serum (FBS) (SIGMA, F2442) and 1x NEAA (SIGMA, M7145); the human leukemia cell line THP-1 (Cytion, 300356) grown in RPMI 1640 (ThermoFisher, catalog no. 11875-093) supplemented with 10% FBS; as well as CHO cells (Cytion, 603479) grown in Ham’s F-12K medium (Gibco, 21127022) supplemented with 10% FBS. Stem cells were grown on Matrigel Matrix (Corning, 35248) in mTeSR1 medium (StemCell Technologies, 05851) with supplement (StemCell Technologies, 05852) that was replaced daily. At _∼_80% confluence, stem cells were dissociated using EDTA (VWR, 437012C) and split 1:6 to 1:10 in medium supplemented with 10 μM Rho-associated protein kinase (ROCK) inhibitor Y-27632 (Calbiochem, 688000) for one day after splitting. For other cells, media was replaced every second day and cells were split 1:6 to 1:10 once per week. All cell lines were authenticated by the supplier via certificate of analysis and additionally in-house by checking morphology. All cell lines were tested negative for mycoplasma contamination before and after the experiments.

### Electroporation

Cells were treated with TrypLE (Gibco, 12605010) for 5 min at 37°C and triturated to obtain single cells, before addition of preheated media. Cells were counted using the Countess Automated Cell Counter (Invitrogen) and cell suspensions were centrifuged at 300 x g for 3 min at room temperature. Cas9 inducible iCRISPR cells were incubated in medium containing 2 μg/ml doxycycline (Clontech, 631311) to express Cas9. For cells without integrated iCRISPR, we used 252 pmol recombinant Streptococcus pyogenes Cas9-HiFi protein from Integrated DNA Technologies or the Cas9 adenine base editor ABE8e (Protein Production Sweden PPS). Electroporation for all cell types was done using the B-16 program of the Nucleofector 2b device (Lonza) in cuvettes for 100 μl Human Stem Cell nucleofection buffer (Lonza, VVPH-5022), containing ∼10^6^ cells, 100Lpmol of electroporation enhancer and 320Lpmol of gRNA or 5 μg of dsDNA for expression of gRNAs where applicable.

### Illumina library preparation and sequencing

Five days or more after transfection cells were dissociated using TrypLE (Gibco, 12605010), pelleted and resuspended in 15 μl of QuickExtract DNA extraction solution (Lucigen, QE09050). Incubation at 65°C for 10 min, 68°C for 5 min and finally 98°C for 5 min was performed to yield single-stranded DNA. PCR was done in a T100 Thermal Cycler (Bio-Rad) using the KAPA2G Robust PCR Kit (Sigma, KK5024) with supplied buffer B and 3 μl of cell extract in a total volume of 25 μl. The thermal cycling was: 95°C for 3 min; 34× (95°C for 15 s, 65°C for 15 s, 72°C for 15 s); 72°C for 60 s. Illumina adapters (P5 and P7) with sample-specific indices were added in a second PCR reaction using Phusion HF MasterMix (Thermo Scientific, F-531L), 0.3 μl of the first PCR product and cycling was: 98°C for 30 s; 25× (98°C for 10 s, 58°C for 10 s, 72°C for 20 s); 72°C for 5 min. Amplifications were analyzed using 2% EX agarose gels (Invitrogen, G4010–11), indexed amplicons were purified using solid phase reversible immobilization beads in a 1:1 ratio of beads to PCR solution^87^.

Double-indexed libraries were sequenced on a MiSeq (Illumina) giving paired-end sequences of 2L×L150Lbp (+7 bp index). After base calling using Bustard (Illumina), adapters were trimmed using leeHom^88^.

### Amplicon sequence analysis

Bam files were demultiplexed and converted into fastq files using SAMtools^89^. Fastq files were used as input for CRISPResso^90^ to analyze sequencing read percentage of wild type (unedited), indels, or targeted nucleotide substitution in case of base editing. Analysis was restricted to amplicons with a minimum of 70% similarity to the wild-type sequence and to a window of 20 bp from each gRNA. Sequence similarity for a targeted nucleotide substitution (HDR command for base editing) occurrence was set to 90%. Unexpected substitutions were ignored as putative sequencing errors.

## DATA AVAILABILITY

RNA-seq datasets have been deposited within NCBI BioProject 1461901 to be made publicly available.

## ACKNOWLEDGEMENTS

We thank the Center for Information Services and High-Performance Computing (ZIH) at TU Dresden for computer time. We acknowledge Protein Production Sweden (PPS) for providing the recombinant ABE8e base editor. PPS is funded by the Swedish Research Council as a national research infrastructure. We thank Dr. Grégory Resch (Laboratory of Bacteriophages and Phage Therapy, Lausanne University Hospital (CHUV), Switzerland) for sharing *Streptococcus tigurinus* 859. This work was supported through the European Research Council (865973 to C.L.B.), the Deutsche Forschungsgemeinschaft SPP 2141 program (468749960 to Z.W., BE 6703/2-1 to C.L.B.), the German Academic Exchange Service (DAAD) (01IM19001 to M.F.), the Foundation Immune Engineering and its Botnar Institute of Immune Engineering (to C.L.B.), Bioprotection Aotearoa (to P.C.F, D.M.M., N.B.), a James Cook Research Fellowship from the Royal Society, Te Apārangi, New Zealand (to P.C.F.), a University of Otago Doctoral Scholarship (to J.L.), a University of Otago Health Sciences Career Development Postdoctoral Fellowship (to N.B.), a New Zealand Health Research Council Explorer Grant (to C.M.B) and a research grant from VILLUM FONDEN (VIL60763 to R.P.-R.). We are grateful to Dr. Rachael E. Workman and Dr. Joshua Modell from Johns Hopkins University for early discussions about orphan tracrRNA elements.

## AUTHOR CONTRIBUTIONS

**Maximilian Feussner**: Conceptualization; Methodology; Formal analysis; Investigation; Writing

- Review & Editing; Visualization; Funding acquisition. **Angela Migur**: Conceptualization; Methodology; Validation; Formal analysis; Investigation; Writing - Review & Editing; Visualization. **Jinquan Li**: Conceptualization; Methodology; Validation; Formal analysis; Investigation; Funding acquisition. **Stephan Riesenberg**: Conceptualization; Methodology; Validation; Formal analysis; Investigation; Writing - Review & Editing; Funding acquisition. **Nils Birkholz**: Methodology; Writing - Review & Editing; Supervision; Funding acquisition. **David Mayo-Muñoz**: Methodology; Writing - Review & Editing; Supervision; Funding acquisition. **Holly Wakelin**: Investigation. **Chris M. Brown**: Conceptualization; Software; Supervision; Funding acquisition. **Zasha Weinberg**: Conceptualization; Software; Writing - Review & Editing; Supervision; Project administration; Funding acquisition. **Rafael Pinilla-Redondo**: Conceptualization; Writing - Original Draft; Writing - Review & Editing; Supervision; Project administration; Funding acquisition. **Peter C. Fineran**: Conceptualization; Writing - Original Draft; Writing - Review & Editing; Supervision; Project administration; Funding acquisition. **Chase L. Beisel**: Conceptualization; Writing - Original Draft; Writing - Review & Editing; Supervision; Project administration; Funding acquisition.

## Supplementary Figures

**Figure S1.**
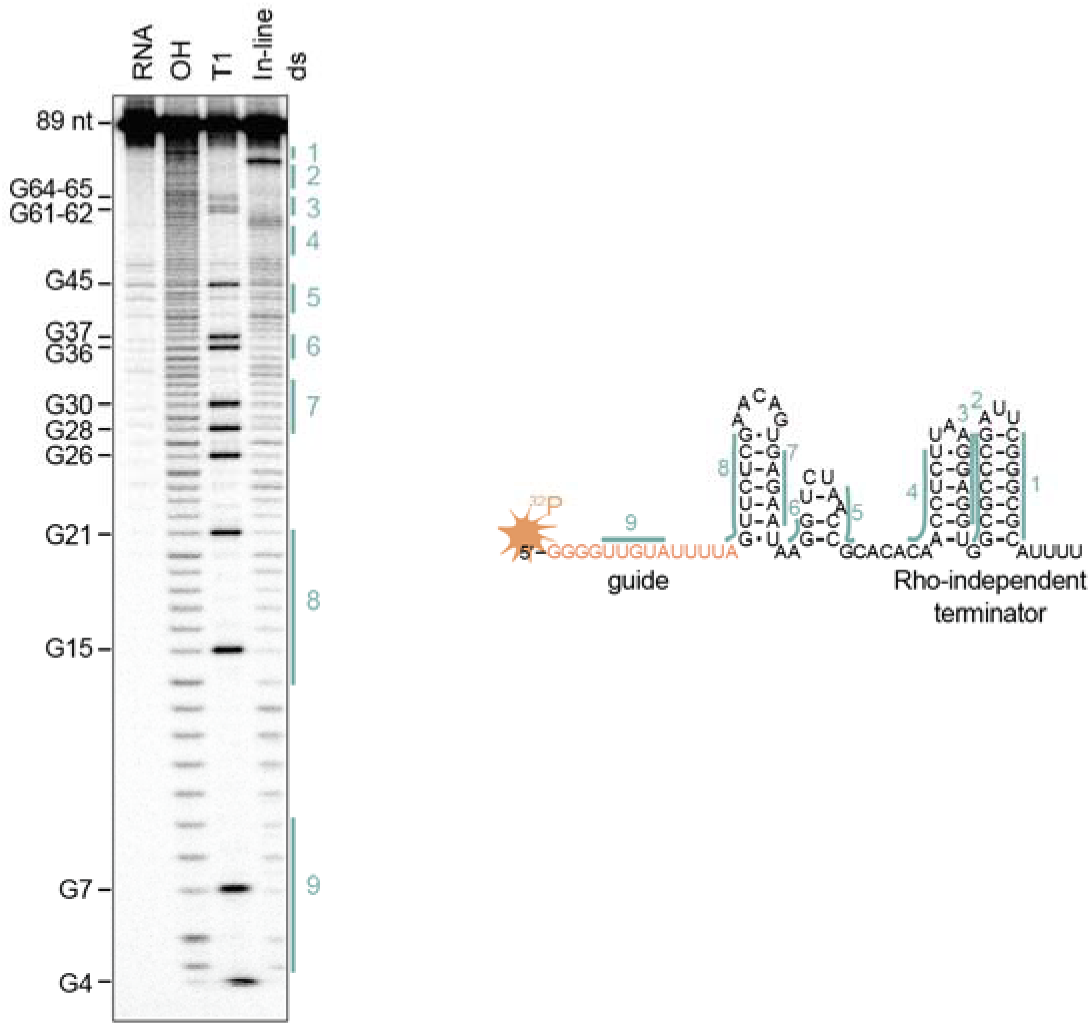
Experimental determination of the *in vitro* structure of the vsgRNA in *S. tigurinus* 859. Left: in-line probing of 5_′_ radiolabeled 89-nt RNA. RNA, untreated. OH, partial alkaline treatment. T1, RNase T1 treatment. Right: predicted minimal-free-energy (MFE) structure of the vsgRNA. The base-paired nucleotides are marked with green bars. As expected, nucleotides predicted to participate in base-pairing show less in-line cleavage than other nucleotides. The gel image is a representative of two independent experiments.

**Figure S2.**
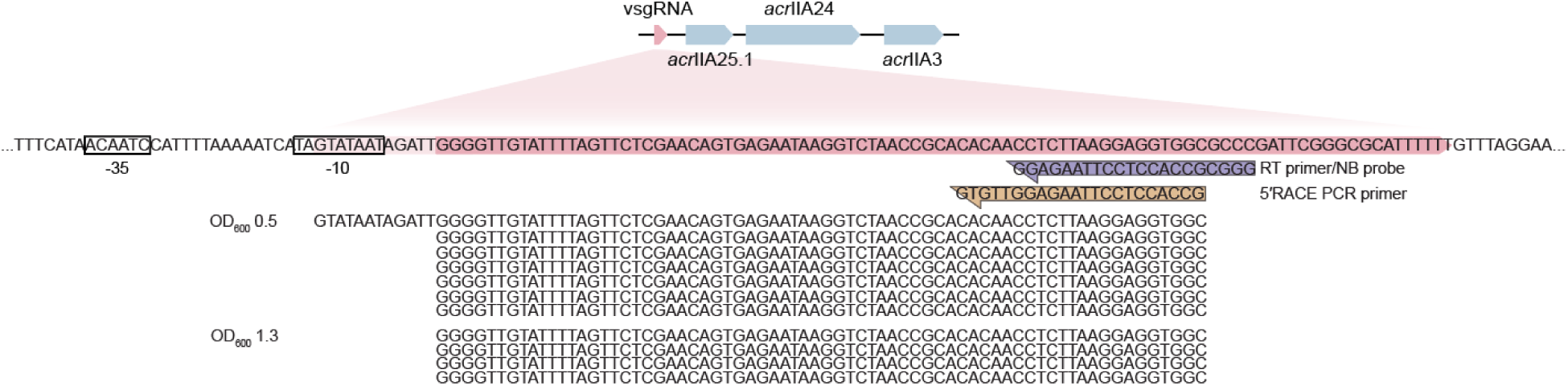
Alignment of the transcripts of the vsgRNA obtained through 5_′_ RACE. The RNA subjected to 5_′_ RACE was extracted from *S. tigurinus* 859 (LNVH01000004.1) cultured to OD_600_ = 0.5 or 1.3. The -10 and -35 elements were predicted using PromoterCalculator (https://www.denovodna.com/software/).

**Figure S3.**
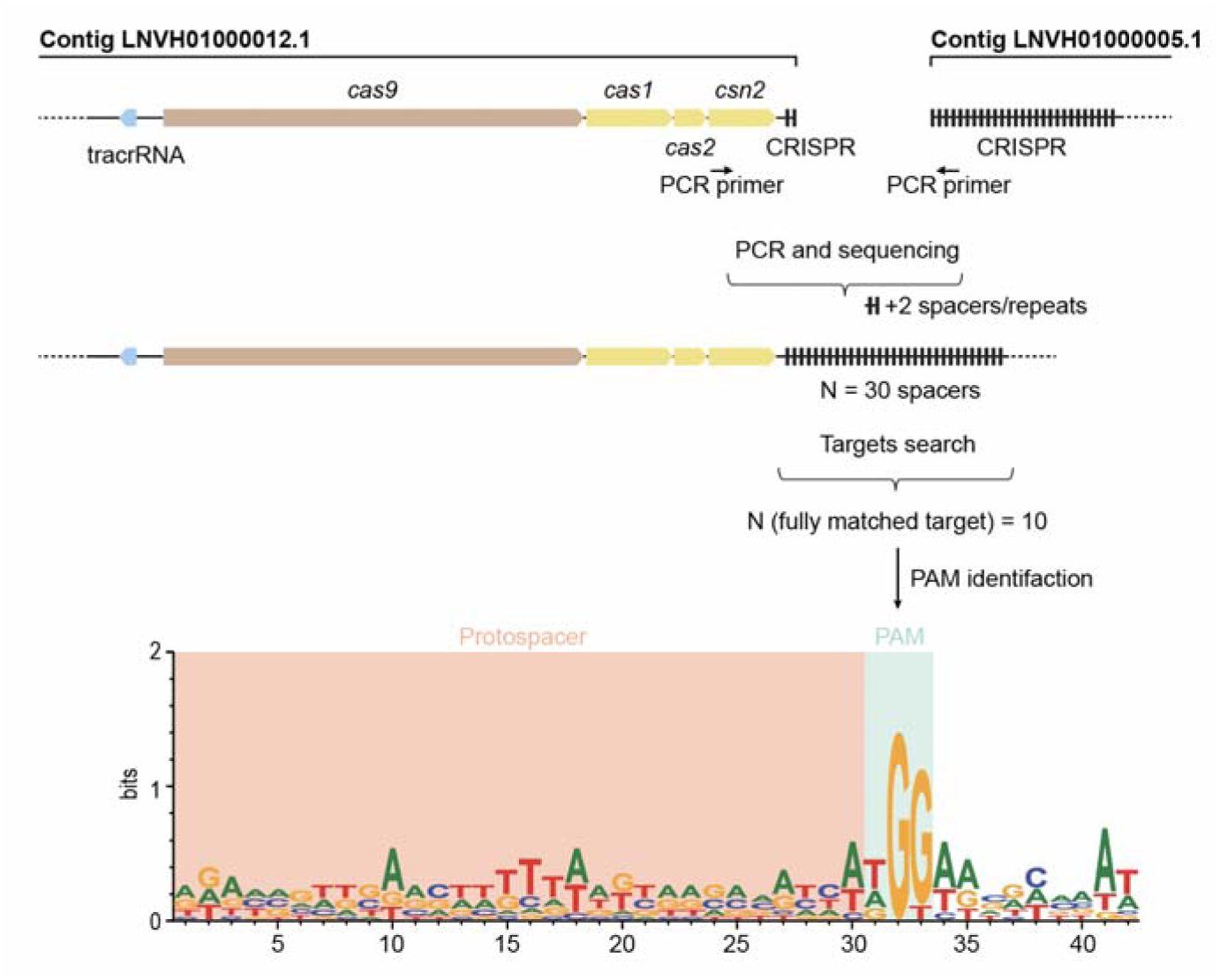
Identification of the PAM sequence of the *S. tigurinus* Cas9. The CRISPR array is located on two different contigs (LNVH01000012.1 and LNVH01000005.1), presumably due to the difficulties of assembling repetitive sequences from short high-throughput-sequencing reads. To find the authentic sequence, we amplified the CRISPR array by PCR and sequenced using Sanger sequencing. The resulting spacer sequences were searched against the NCBI nr nucleotide database, which does not include the *S. tigurinus* 859 genome. Full length matches and their neighbouring sequences were aligned and analyzed to identify the PAM sequence (NGG).

**Figure S4.**
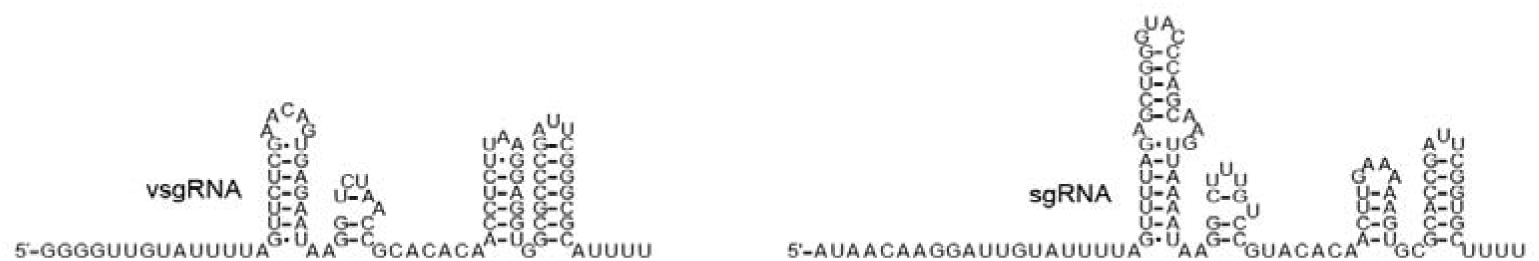
Sequences and RNA secondary structures of the vsgRNA (left) and sgRNA (right) used in the *in vitro* SpCas9 binding assay and in the DNA binding and cleavage assays.

**Figure S5.**
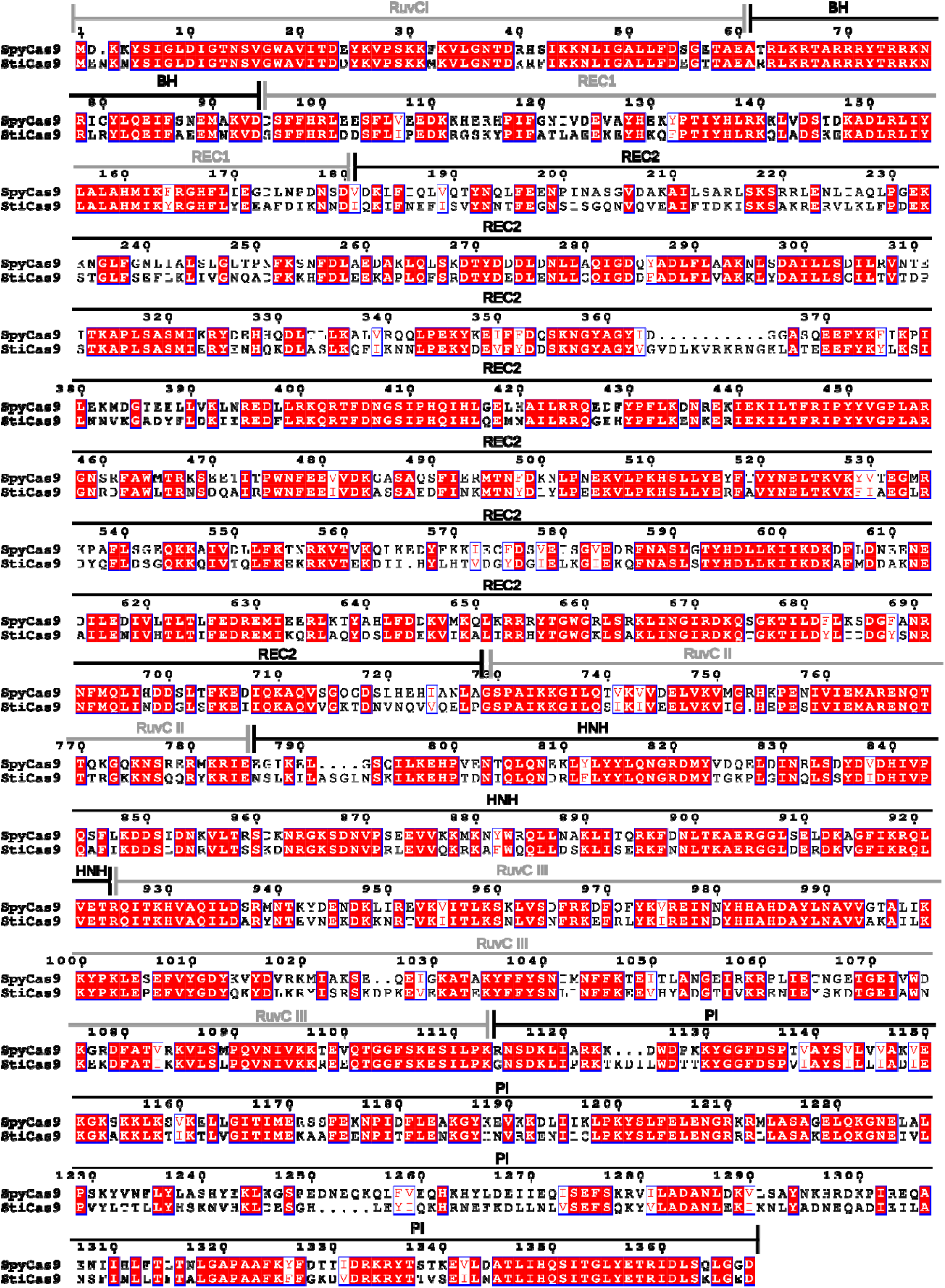
Sequence alignment between *S. tigurinus* Cas9 and *S. pyogenes* Cas9. Protein domains are based on *S. pyogenes* Cas9 and are visualized above the amino acid sequences (https://espript.ibcp.fr/ESPript/ESPript/). Conserved amino acids are highlighted in red.

**Figure S6.**
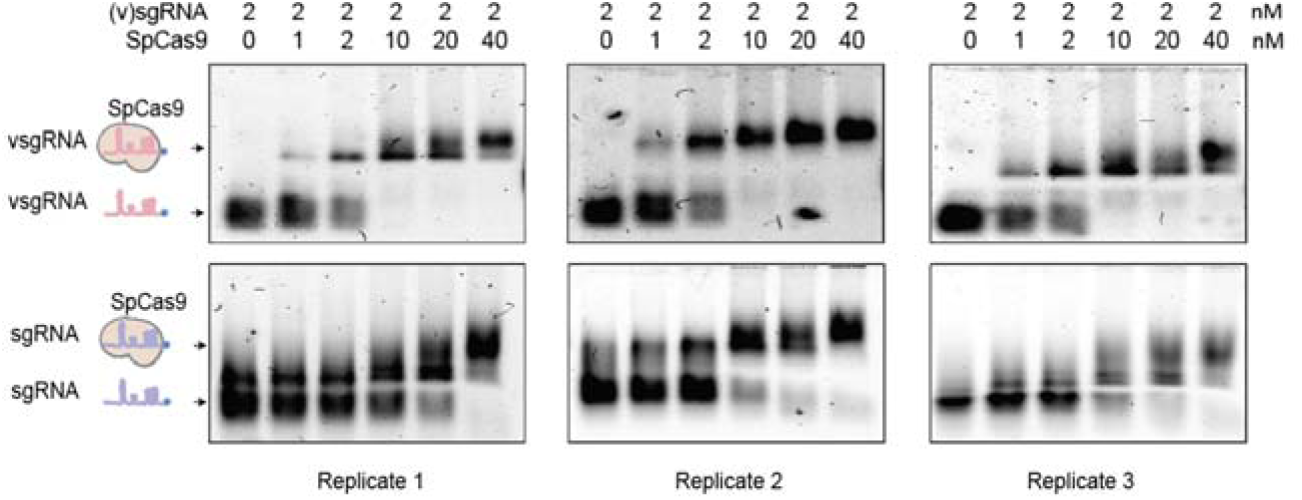
Individual EMSAs assessing vsgRNA and sgRNA binding to SpCas9. The bands running at the size of the RNA alone were used to calculate the percentage of the unbound RNA.

**Figure S7.**
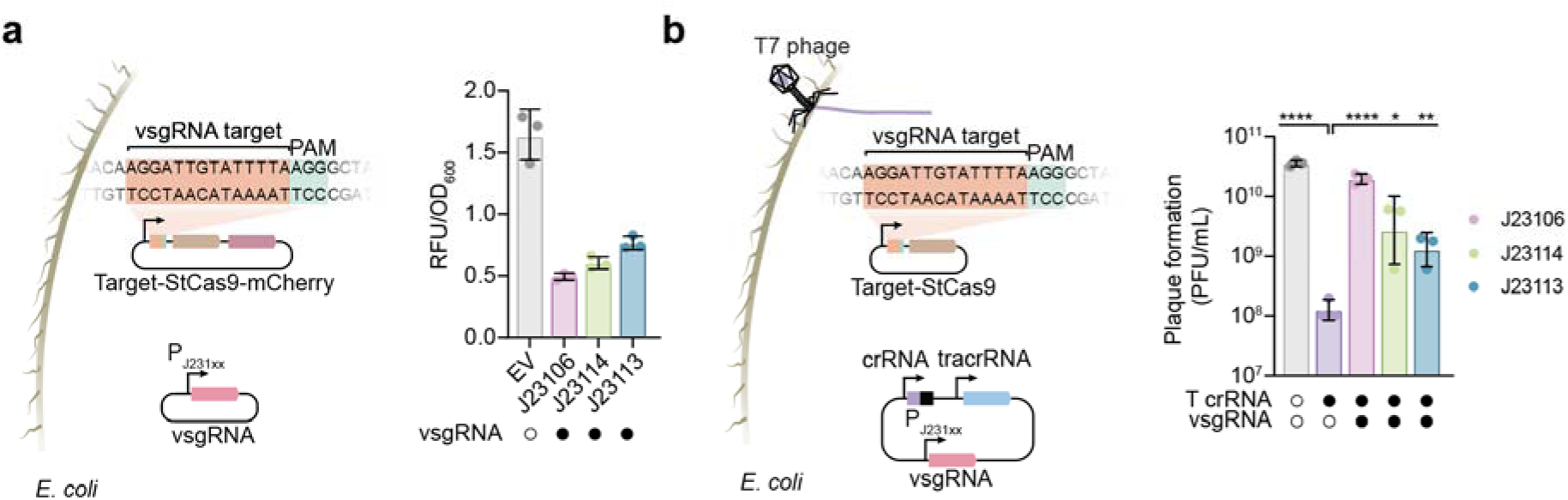
Dose-dependent promoter targeting by vsgRNA. **a,** PAM-dependent silencing of a mCherry reporter by the vsgRNA in *E. coli*. EV, empty vector. RFU, Relative Fluorescence Units. vsgRNA was expressed from three different constitutive promoters ranging in activity from J23106 (strongest) to J23113 (weakest) (https://parts.igem.org/Promoters/Catalog/Anderson). Circles represent individual measurements. Bars and error bars represent the geometric mean and geometric standard deviation from three biological replicates. **b,** Impact of promoter targeting with the vsgRNA on Cas9-mediated immunity against infection by T7 phage. T, targeting. Cas9 was expressed from a constitutive promoter followed by the vsgRNA target site. vsgRNA was expressed from three different constitutive promoters as in **Figure S7a**. Circles represent individual measurements. Bars and error bars represent the geometric mean and geometric standard deviation from three biological replicates. *, p < 0.05. **, p < 0.01. ***, p < 0.001. ****, p < 0.0001 based on an unpaired t-test on geometric means with two tails.

**Figure S8.**
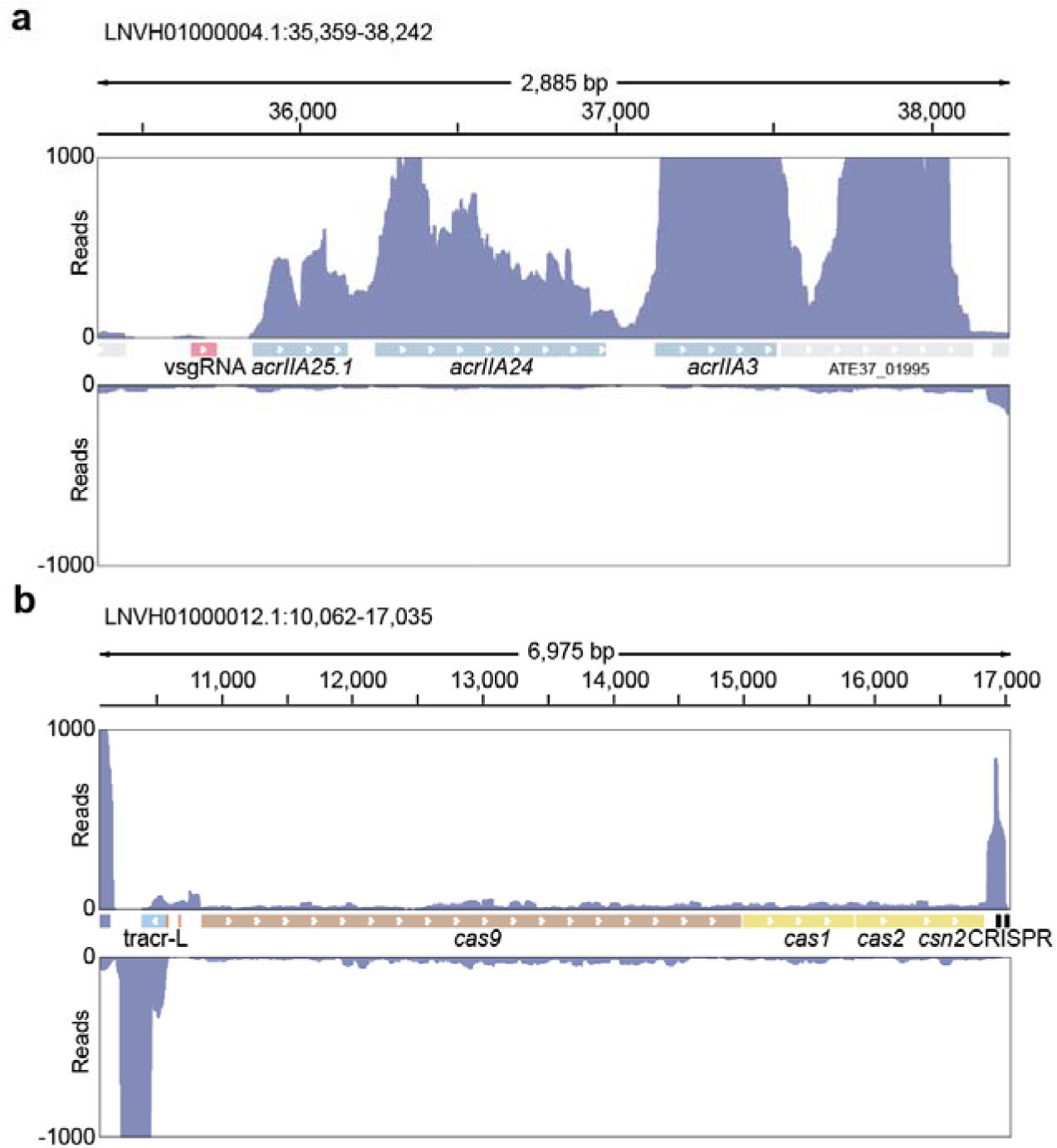
RNA-seq analysis of the prophage and *cas9* loci in *Streptococcus tigurinus* 859. **a,** RNA-seq reads from the total RNA of *S. tigurinus* mapped to the prophage locus with vsgRNA, *acrIIA25.1*, *acrIIA24*, and *acrIIA3* (LNVH01000004.1). The low read count for the vsgRNA can be attributed to RNA fragmentation as part of the NGS library preparation. One representative biological replicate out of three is shown. **b,** RNA-seq reads from the small size fraction of total RNA of *S. tigurinus* mapped to the *cas9* locus (LNVH01000012.1). One representative biological replicate out of three is shown.

**Figure S9.**
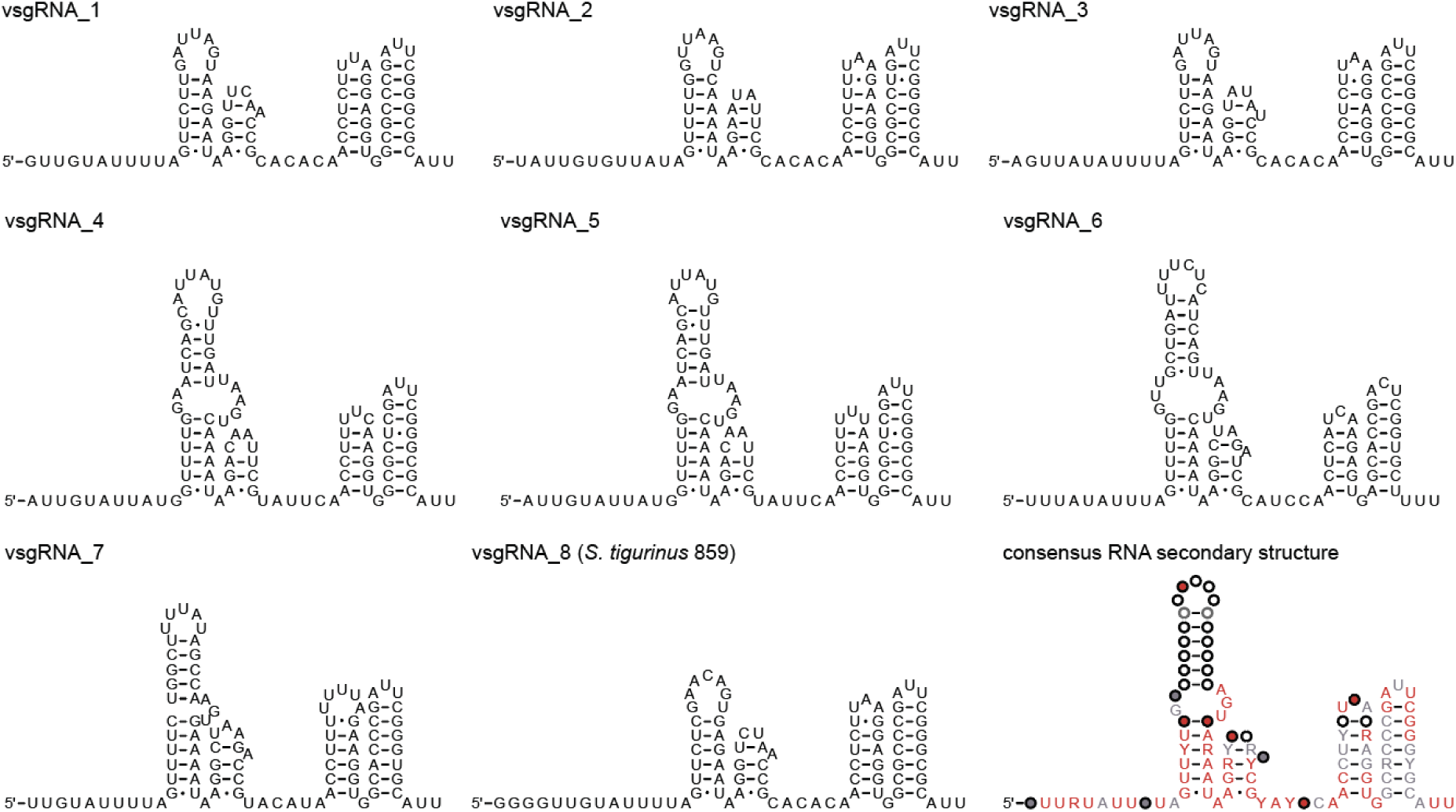
Sequence and RNA secondary structures of experimentally tested vsgRNAs with their guide sequences. RNA secondary structure is based on the covariance models used to identify the vsgRNA. The consensus RNA secondary structure is based on a multiple sequence alignment of all eighth vsgRNA and matches that shown in Figure 4c. Red nucleotides indicate those present in all eighth vsgRNA at this position, gray nucleotides indicate that these nucleotides are present in at least six vsgRNAs. Different colored circles represent no sequence conservation at this position. Nucleotide accession numbers and the position of the vsgRNA can be found in **Table S3.**

**Figure S10.**
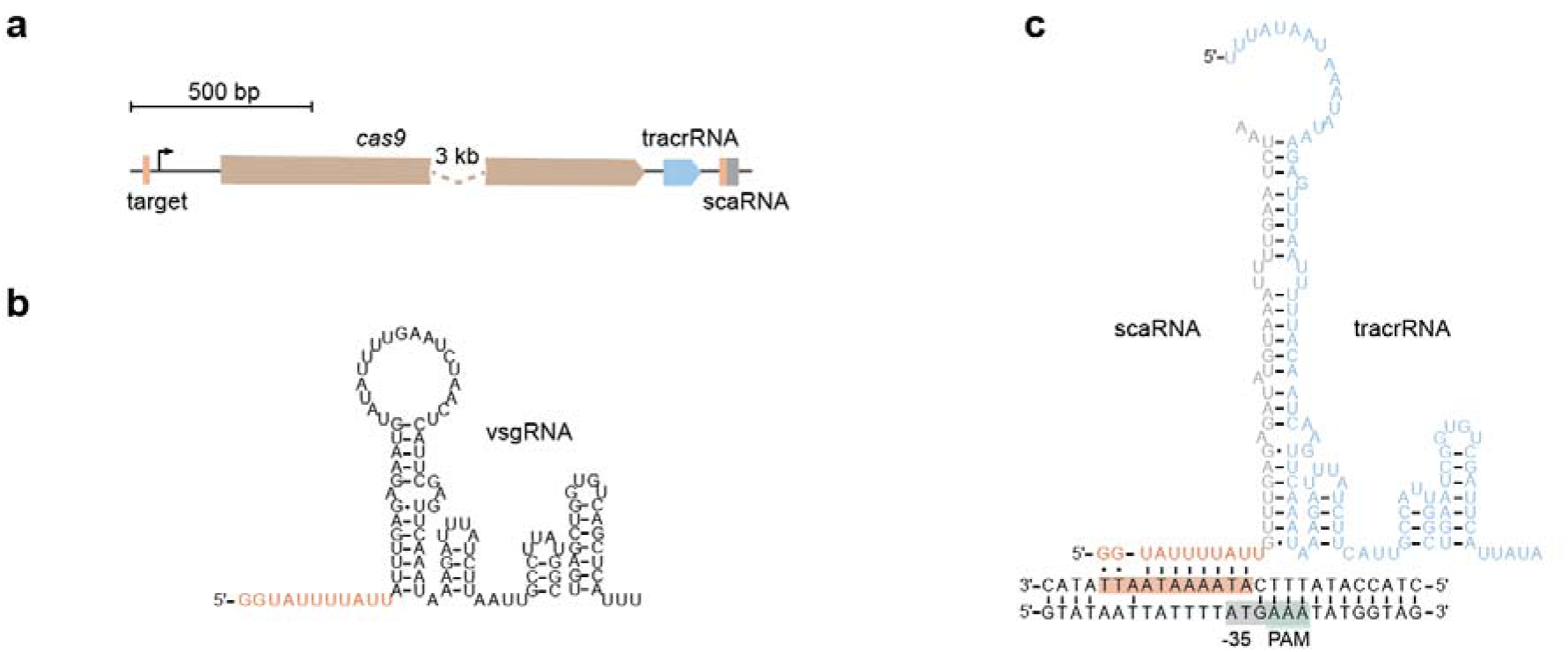
The identified vsgRNA and scaRNA in *Gemella morbillorum* (NZ_CAURBT010000002.1). **a,** The *cas9* and scaRNA locus in *Gemella*. The scaRNA guide and its DNA target are in orange. The system lacks an identifiable CRISPR array. **b,** Sequence and RNA secondary structure of the vsgRNA. **c,** Predicted RNA secondary structure of the scaRNA-tracrRNA duplex and the DNA target within the *cas9* promoter. The PAM sequence was predicted with Protein2PAM^30^ using the Cas9 sequence.

**Figure S11.**
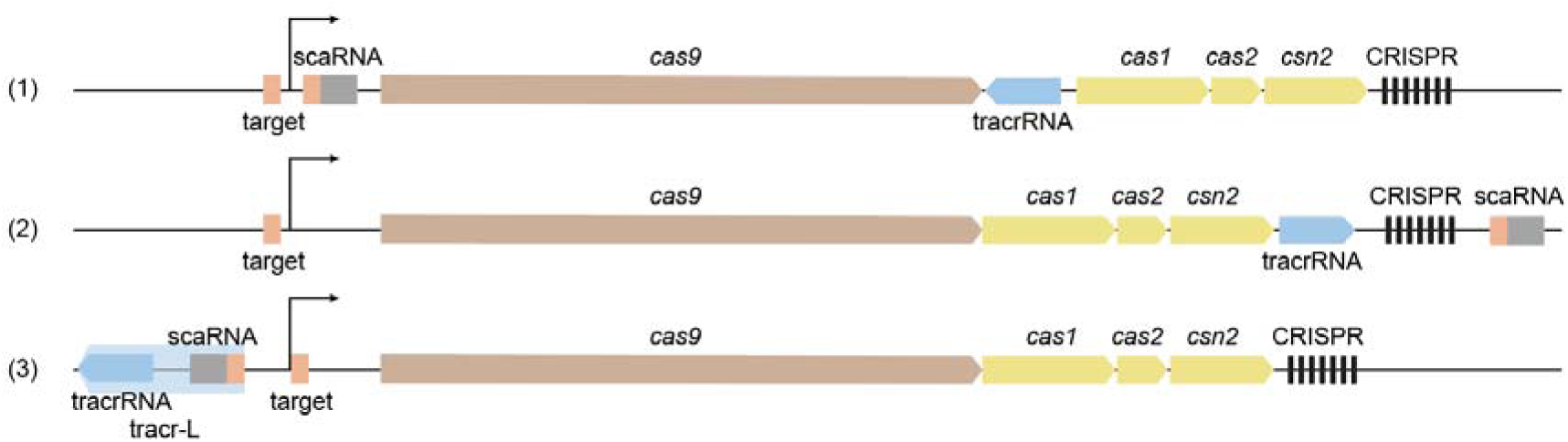
Genomic locations of the scaRNA suggest the origin of tracr-L. The scaRNA consists of a repeat (gray) and guide (orange) and can be found in three locations: (1) within the 5_′_ UTR of *cas9*, (2) downstream of the CRISPR array, or (3) upstream of the tracrRNA. For the last case, the scaRNA is cotranscribed with the tracrRNA, forming tracr-L. The scaRNA/tracr-L target site (also orange) is near the *cas9* transcriptional start site.

**Figure S12.**
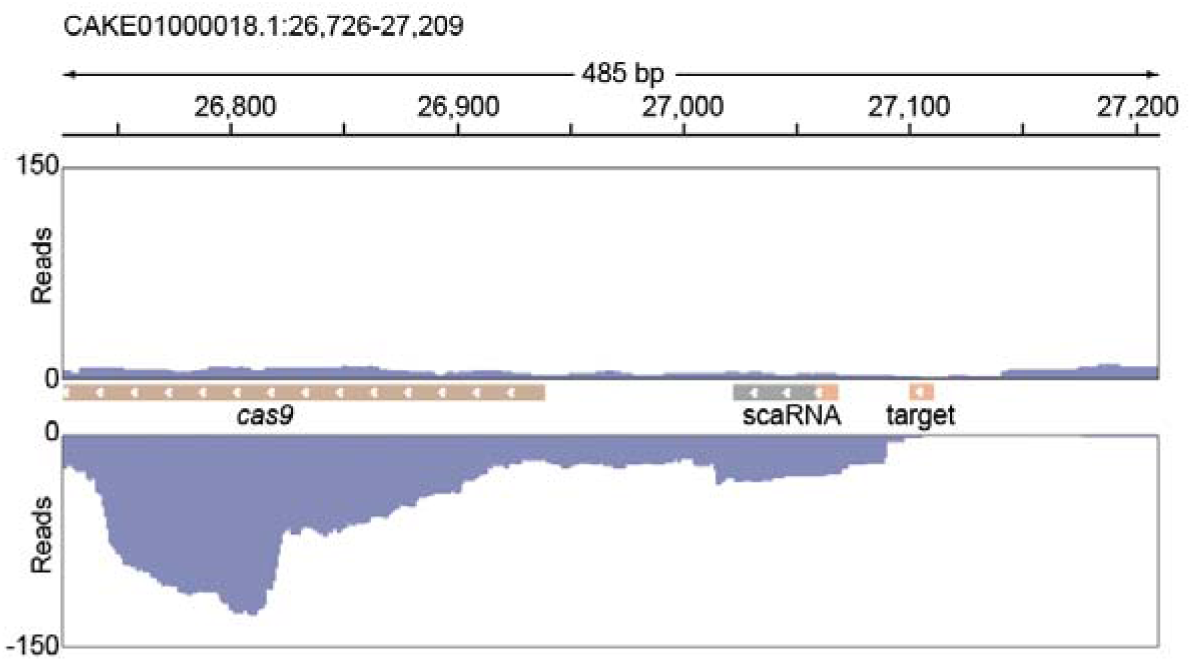
RNA-seq reads from the enriched small fraction of total RNA without fragmentation of *L. hominis* mapped to scaRNA-*cas9* locus (CAKE01000018.1). One of three representative biological replicates is shown. The scaRNA, consisting of a repeat (gray) and guide (orange), is co-transcribed with *cas9* (brown). The scaRNA target site (also orange) is immediately upstream of the shared transcriptional start site.

**Figure S13.**
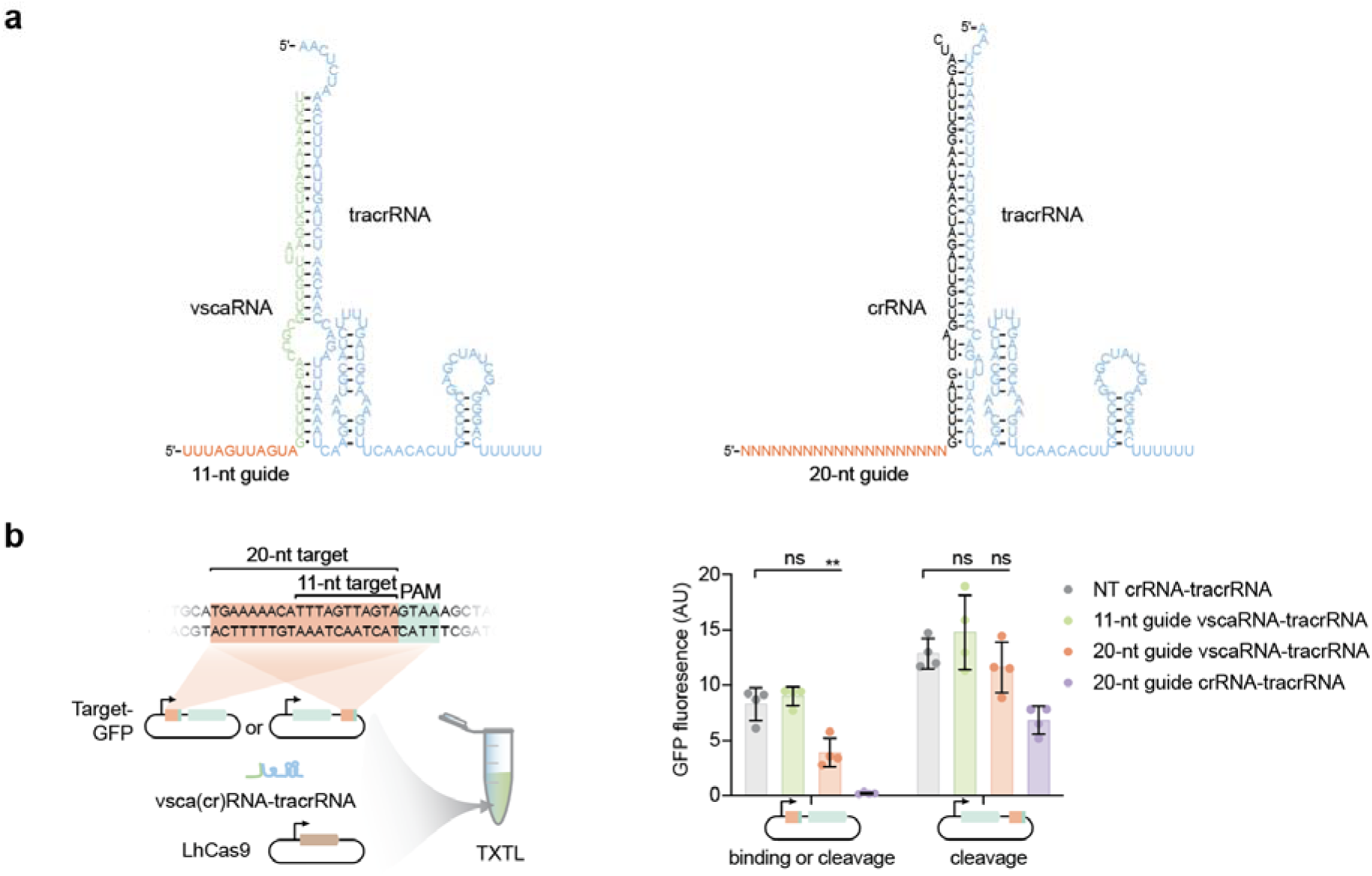
Gene repression by the Cas9-regulating vscaRNA found in *Lactobacillus gasseri* (NZ_JARPOE010000001.1 and NZ_JAQOUF010000001.1). **a,** Sequence and RNA secondary structure of the vscaRNA (left) or crRNA (right) paired to the tracrRNA from *Lactobacillus hominis* DSM 23910. **b,** Left: overview of the combined components in a TXTL reaction. vsca(cr)RNA-tracrRNA indicates that the reaction contains tracrRNA with either vscaRNA or crRNA. Right: GFP fluorescence following DNA targeting with different guide RNAs. NT, non-targeting. The 11- or 20-nt guides correspond to the 11- or 20-nt target, respectively. Circles represent individual measurements. Bars and error bars represent the mean and standard deviation from four independent measurements. *, p < 0.05. **, p < 0.01 based on an unpaired t-test with two tails.

**Figure S14.**
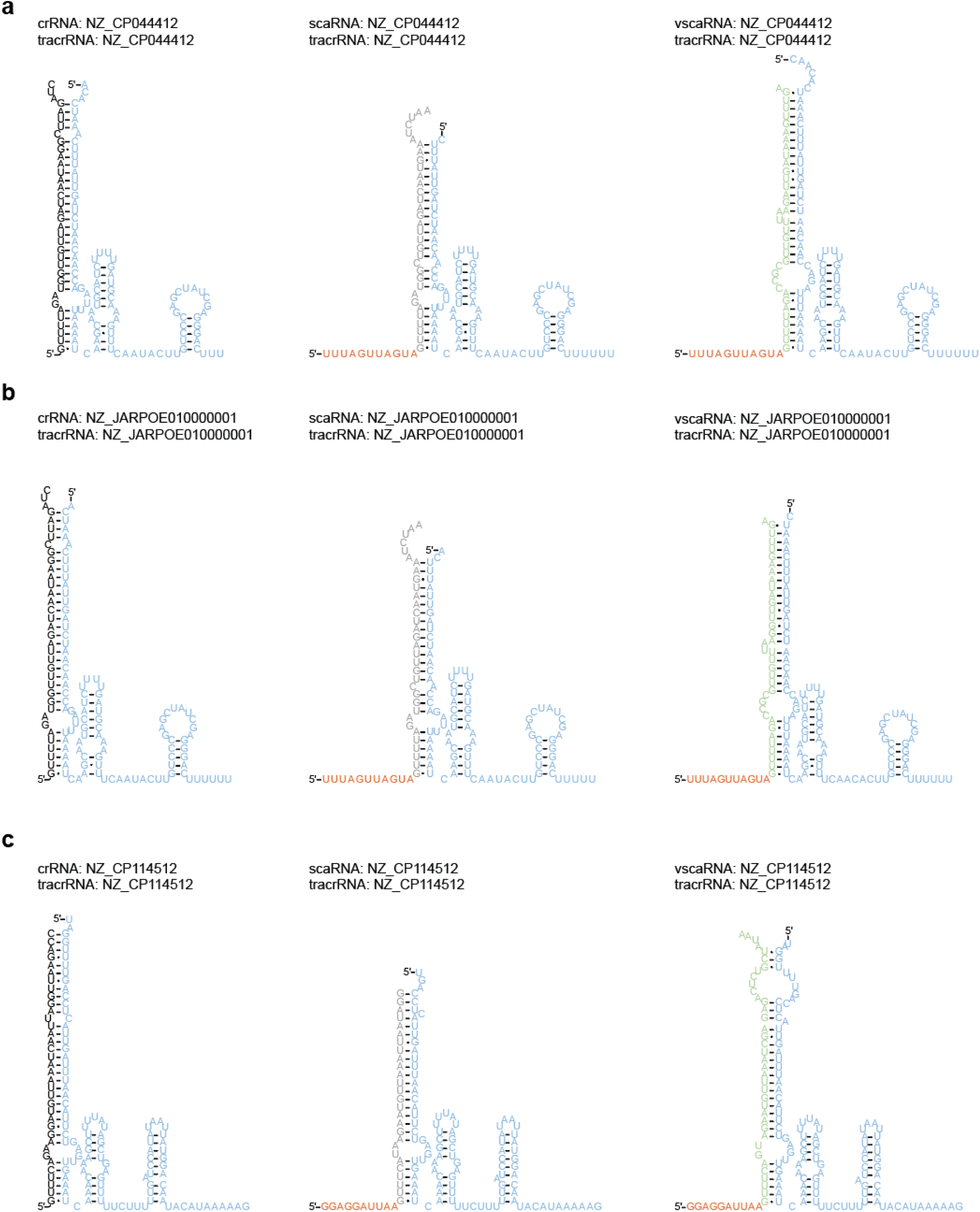
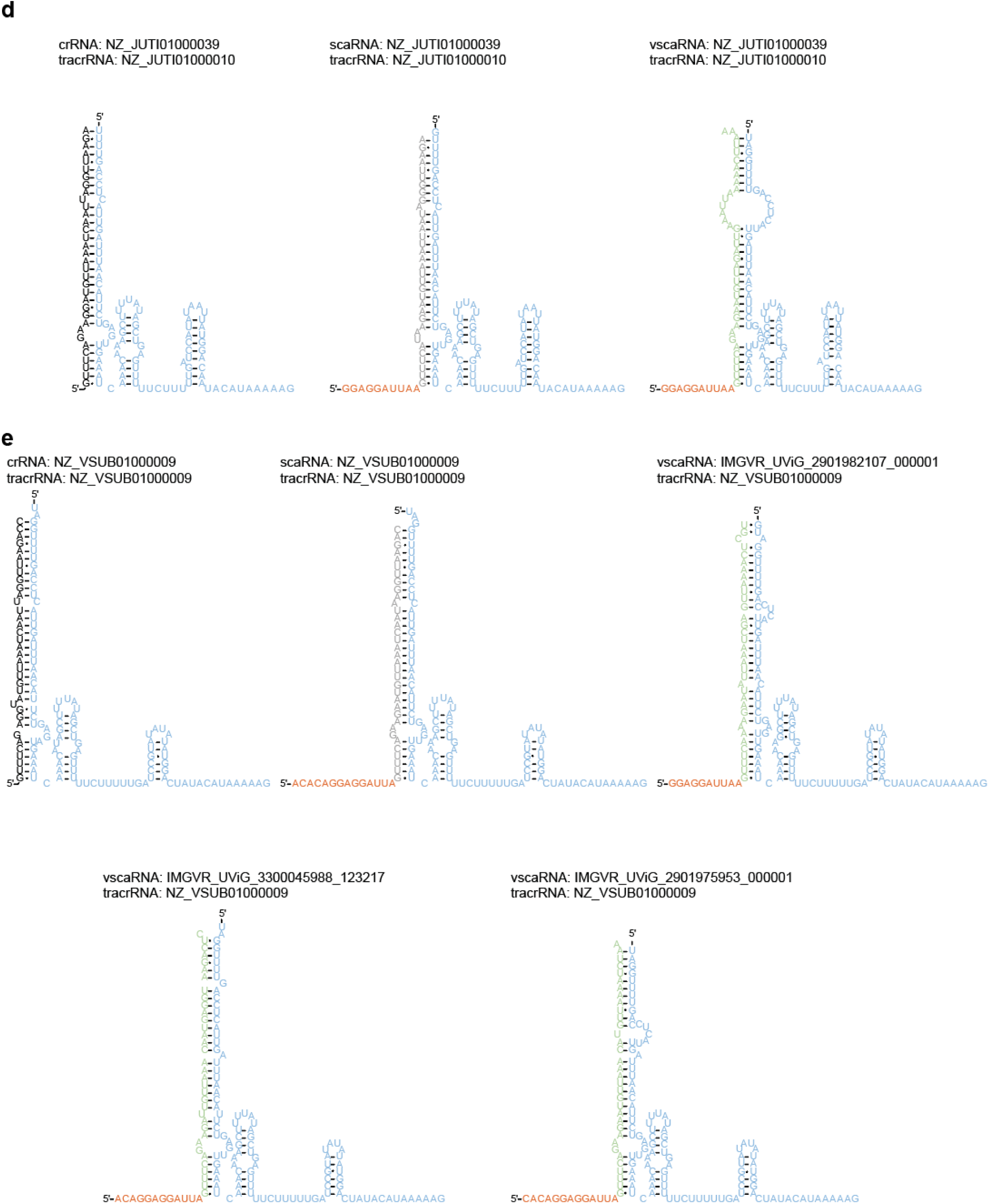
crRNA-tracrRNA, scaRNA-tracrRNA, and vscaRNA-tracrRNA pairs: sequences and RNA secondary structures. The accession number of the genome in which each RNA was identified is shown above the corresponding structure. The guide of the scaRNA or vscaRNA is orange, the crRNA repeat is black, the crRNA repeat-like element of the scaRNA or vscaRNA is gray or green, and the tracrRNA is blue. **a,** Pairs identified in *Lactobacillus sp.* JM1 (NZ_CP044412). **b,** Pairs identified in *Lactobacillus gasseri* strain IM13 (NZ_JARPOE010000001.1). **c,** Pairs identified in *Ligilactobacillus salivarius* strain S39 (NZ_CP114512.1). **d,** Pairs identified in *Ligilactobacillus salivarius* strain 778 (NZ_JUTI00000000.1). **e,** crRNA-tracrRNA and scaRNA-tracrRNA pairs identified in *Ligilactobacillus salivarius* strain FYNDL5_1 (NZ_VSUB01000009). vscaRNAs were identified in the IMG/VR database and paired to the tracrRNA from NZ_VSUB01000009 genome.

**Figure S15.**
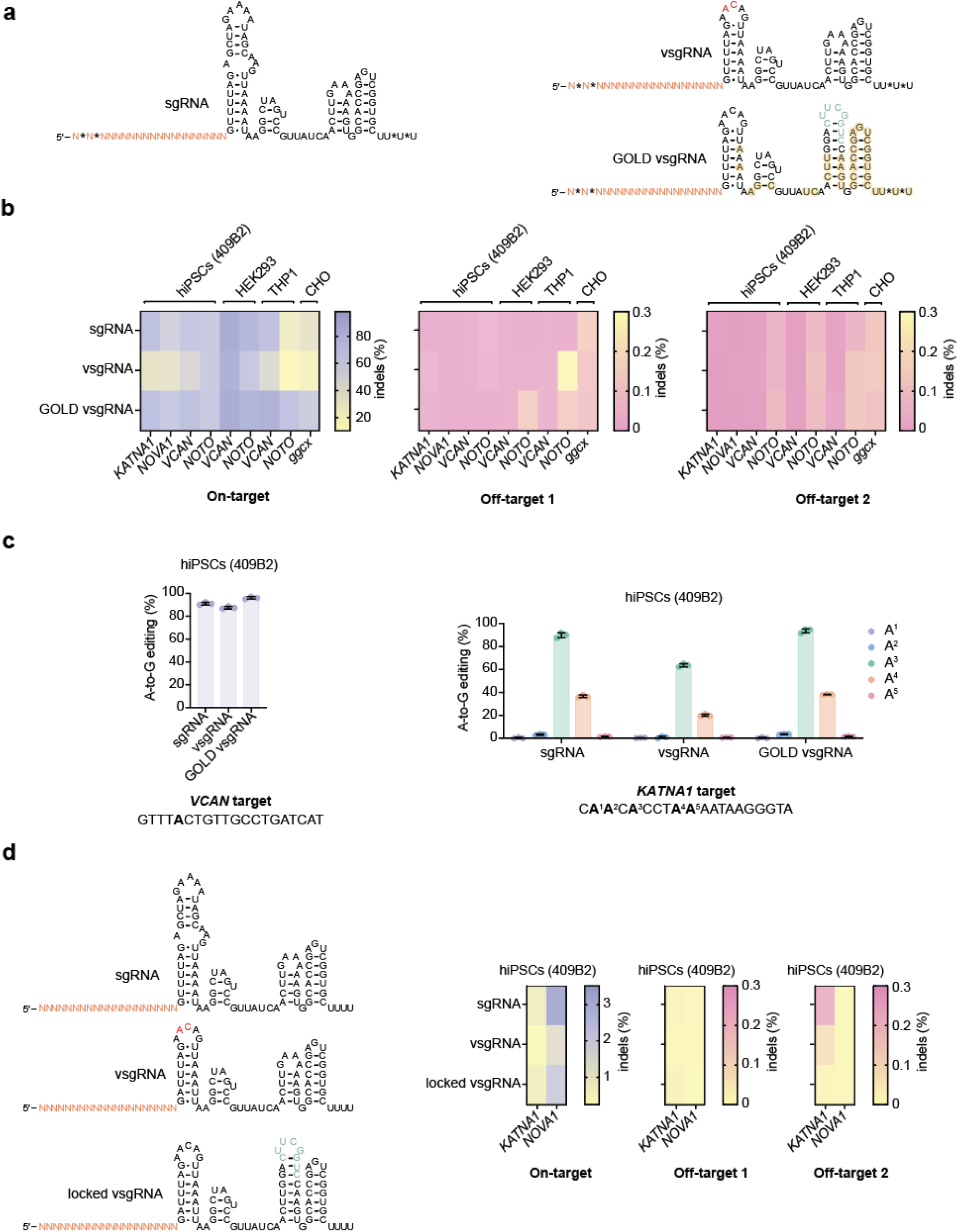
Modified vsgRNAs enact gene editing in human cells. **a,** Comparison of the sgRNA, vsgRNA, and GOLD vsgRNA sequences and RNA secondary structures. The nucleotides that differ in sgRNA and vsgRNA are in red. The nucleotides that differ in vsgRNA and GOLD vsgRNA are in green. 20 nt guide is in orange. *, phosphorothioate bond. The nucleotides with 2′-OMe modifications are circled with yellow. **b,** Indel formation across five tested sites in four cell lines. Each color in the heat map represents the mean of three independent measurements. Off-target indel formation for each gRNA was assessed at two different sites. **c,** A-to-G base editing in hiPSCs (409B2) was evaluated at two target sites, with the edited adenosine residues indicated in bold. **d,** Indel formation at two challenging editing sites when gRNAs are expressed from a U6 promoter in an induced pluripotent cell line. Left: Comparison of the sgRNA, vsgRNA, and locked vsgRNA sequences and RNA secondary structures. The nucleotides that differ in sgRNA and vsgRNA are in red. The nucleotides that differ in vsgRNA and locked vsgRNA are in green. The 20-nt guide is in orange. Right: indel formation results. Each color in the heat map represents the mean of three independent measurements. Off-target indel formation for each gRNA was assessed at two sites.

## SUPPLEMENTARY TABLES

**Table S1.** sgRNAs used to build co-variance models.

**Table S2.** Predicted vsgRNA sequences.

**Table S3.** Experimentally verified vsgRNA sequences.

**Table S4.** Predicted tracr-L sequences.

**Table S5.** Predicted scaRNA sequences.

**Table S6.** Predicted vscaRNA sequences.

**Table S7.** Strains used in this work.

**Table S8.** DNA oligos used in this work.

**Table S9.** Plasmids used in this work.

**Table S10.** RNA sequences used in this work.

**Table S11.** gBlocks used in this work.

All supplementary tables are contained as tabs within the provided Excel spreadsheet.

